# Molecular Parallelisms and Divergences Between Human and Canine Cancers Would a dog be an appropriate experimental model for human cancers?

**DOI:** 10.1101/2020.10.01.318188

**Authors:** Sadaf Ambreen, Li Guanhan, Binbin Zhao, Gang Bao, Yiwen Song, Degui Lin, Dafei Wu, Yali Hou, Xuemei Lu

**Author notes:** These authors contributed equally to this work.

## Abstract

Canine mammary cancer is poorly characterized at the genomic level. Dog really can be an appropriate experimental model for human cancers from the genomic and evolutionary perspective or not? Here, we perform a cross-species cancer genomics analysis, independent evolution of cancer from normal tissues, which provide us an excellent opportunity to address an evolutionary perspective of cancer. As, evolutionary theories are critical for understanding tumorigenesis at the level of species as well as at the level of cells and tissues, for the development of effective therapies. Analysis of canine mammary cancer reveals a diversity of histological types as compare to human breast cancer. Our systematic analysis of 24 canine mammary tumors with whole-genome sequences, reveals 185 protein-coding cancer genes carried exonic mutations. Cross-species comparative analysis of 1080 human breast cancers identifies higher median mutation frequency in human breast cancer and canine mammary cancer shows lower across exonic regions (2.67 and 0.187 average no. of mutations per tumor per megabase (Mb), respectively). A comparison of somatic mutations in the *PIK3CA* gene, reveals common recurrence of the conserved mutations, in both species. However, the Ka/Ks ratio in the human *PIK3CA* gene 2.37 is higher and 1.43 in dogs is lower. To address the mutation accumulation and antagonistic pleiotropy theory, we investigated Ka/Ks value 237 aging-related genes from human and canine, the aging-related genes do not show selection in canine mammary cancer. It demonstrates new aspects of cancer genes that are evolving in different species instantaneously. These findings may suggest, the same organs in different mammals impose different selective pressures on the same set of genes in cancer. In both species, some genes may experience strong selective pressures, but do not converge genetically or the conserved genes do not show the same selection pressure in both species. However, human breast cancer shows transcriptomic similarity with canine mammary carcinoma but the other subtypes are quite different. We found canine mammary tumor can be used as a model for inter and intra-tumor heterogeneity. These findings provided insight into mammary cancer across species and possessed potential clinical significance. Collectively, these studies suggest a convergence of some genetic changes in mammary cancer between species but also distinctly different paths to tumorigenesis.

## Introduction

Tumorigenesis has been widely accepted as an evolutionary process that comprises two stages of evolution between tumors and normal tissues (Stage I) and within tumors (Stage II) ^1^. Patterns of mutation and natural selection, the predominant evolutionary driver forces, vary at the two stages based on the evidence of low genetic convergence among different cancer cases revealed by The Cancer Genome Atlas (TCGA) data and of extremely high intra-tumor genetic diversity measured in high-density sampling studies ^2,3^. At Stage, I, positive and negative selection may both exist but neatly counteract in absence of recombination, presenting a plausible neutrality 1, whereas non-Darwinian (neutral) selection was increasingly supported at Stage II by the high-density sampling studies ^2,3^ and comparatively genomic and transcriptional distances among distinct normal and cancerous cell populations ^4^. Deciphering the evolutionary patterns during tumorigenesis such as selectivity or neutrality, adaptive convergence, or divergence is of both theoretical and clinical significance ^5^. Cross-species cancer genomics, independent evolution from normal tissues, provide an excellent opportunity to address this long-standing issue: Does selection drive cancer evolution along with a relatively deterministic (selectivity) or contingent (neutrality) way across species? If there is the selection at Stage I, whether the same organs in different mammals impose similar selection pressures on the same set of genes? Is neutral selection at Stage II supported in different species? If there exists adaptive divergence among species, what are the causes and molecular mechanisms ^6^?

The dog has experienced increased amounts of genetic diseases including cancers subject to human survival and aesthetic preferences, which is widely acknowledged as an increasingly powerful animal model to reveal the genomic signatures of cancer development and progression based on its desired characteristics ^7^. Analogous to human, canine cancers are generally spontaneous and do not need to be induced by exogenous materials like murine cancer models. The spontaneous tumors in dogs share a wide variety of epidemiologic, biologic, and clinical features with human cancer ^8^. Dogs are diagnosed with many of the same cancers as humans, with a similar prevalence of cancer types with few exceptions, and with a similar presentation, pathology, and treatment response ^8^. Cancer biology studies using gene expression data in both human and dog indicate that the target genes and molecular signaling pathways involved in canine cancer development significantly resemble those found in human ^9^, supporting the use of canine cancer model as a homolog for what occurs in physiopathological aspects of cancer biology in human. Like other cancer types, concerning genetic, biological, anatomic, and clinical similarities are documented between canine mammary cancer and human breast cancer. Nevertheless, would dog can be an appropriate experimental model for human cancers from the genomic and evolutionary views?

It is acknowledged that cancer is a significant aging-related disease in both humans and dogs. Most canine cancers start to develop in later life with a sharp increase after the age of 6 years old and a peak at 10, which is similar to human ^10^. Understanding the evolutionary origin of aging-related diseases like cancer and the underlying theory of aging evolution remains largely elusive. Mutation accumulation^11^ and antagonistic pleiotropy^12^ are two prevailing theories proposed to address aging evolution Mutation accumulation theory states that the effects of natural selection decrease as age increases, resulting in maladaptive detrimental mutations that accumulate late in life. Antagonistic pleiotropy theory hypothesizes a trade-off that evolutionary adaptations benefiting the fitness in early life increase disease burden in later life after reproduction ^13^. Both theories haven’t been well verified and exhibit setbacks. Increased incidence of cancer in both humans and dogs provides an opportunity to investigate whether there exits evolutionary similarity or divergence between human and dog cancer, if any, is the similarity attributed to mutation accumulation or antagonistic pleiotropy, and is the divergence related to lineage-specific selective regime?^12^

In this study, we take breast cancer as an example to address the issues of adaptive convergence or divergence in distinct species, because breast cancer is the most common malignancy in women and is the third most common tumor in dogs. We performed whole-genome sequencing in 27 dog mammary gland cancers and reanalyzed 560 WGS and 1080 WES human breast cancers in comparison with their normal samples, profiling the genomic signatures of tumor development and progression, and implemented the comparison of cross-species cancer genomics.

## RESULTS

### Characterization of somatic mutations for canine malignant mammary cancers

To address cross-species cancer genomics questions, we deeply sequenced whole-genomes of 27 malignant mammary tumors from 17 dogs that comprise 8 breeds with an average age of diagnosis of 12, along with the matched normal genomes from the same individuals. Cancers cover varieties of histopathologic classifications, consisting of carcinoma (simple, solid, complex, special types (squamous, adenosquamous, tubulopapillary, lipid)), sarcoma (osteosarcoma, rhabdosarcoma) and carcinosarcoma according to the World Health Organization (WHO) criteria, and various grades according to the Elston and Ellis scoring system, as presented in Fig.1 and Table S1, which exhibit substantially morphological and biological heterogeneity compared with Human breast cancer. Cancers were also classified as subtypes of luminal A, luminal B, human epidermal growth factor receptor 2 (*HER2*)-overexpressed, and triple-negative using immunohistochemistry based on presence of estrogen receptors (*ER*), progesterone receptors (*PR*), and *HER2*.

We performed single-nucleotide variation (SNV) calling in the WGS data from the canine mammary cancers (see Materials and methods). A total of 47,715 somatic single nucleotide variants (SNVs) comprising 180 exonic mutations (97 nonsynonymous, 69 synonymous and 11 stop gain) (Table S4) were identified in the canine mammary neoplasms, with substantial variations in number between and within histologic cancer types (Table S1, Figure 1). Although complex carcinoma predisposes to carry fewest mutations whereas osteosarcoma exhibits the highest, the difference is insignificant based on analysis of variance. 179 genes were identified with exonic mutations in at least one sample, significantly regulating the biological functions of protein kinase activity and cellular chloride ion homeostasis, as well as the pathways in cancer like *PI3K*-*Akt* signaling (*P*<0.05, Table S4, S5). Only 4 genes (*PIK3CA, NEB, PCLO*, and *CKB*) recurrently mutated in at least 2 samples, with a concurrency of 33.3%, 11.11%, 7.4%, respectively, which may involve mammary cancer development in dog (Table S3).

**Figure 1.**
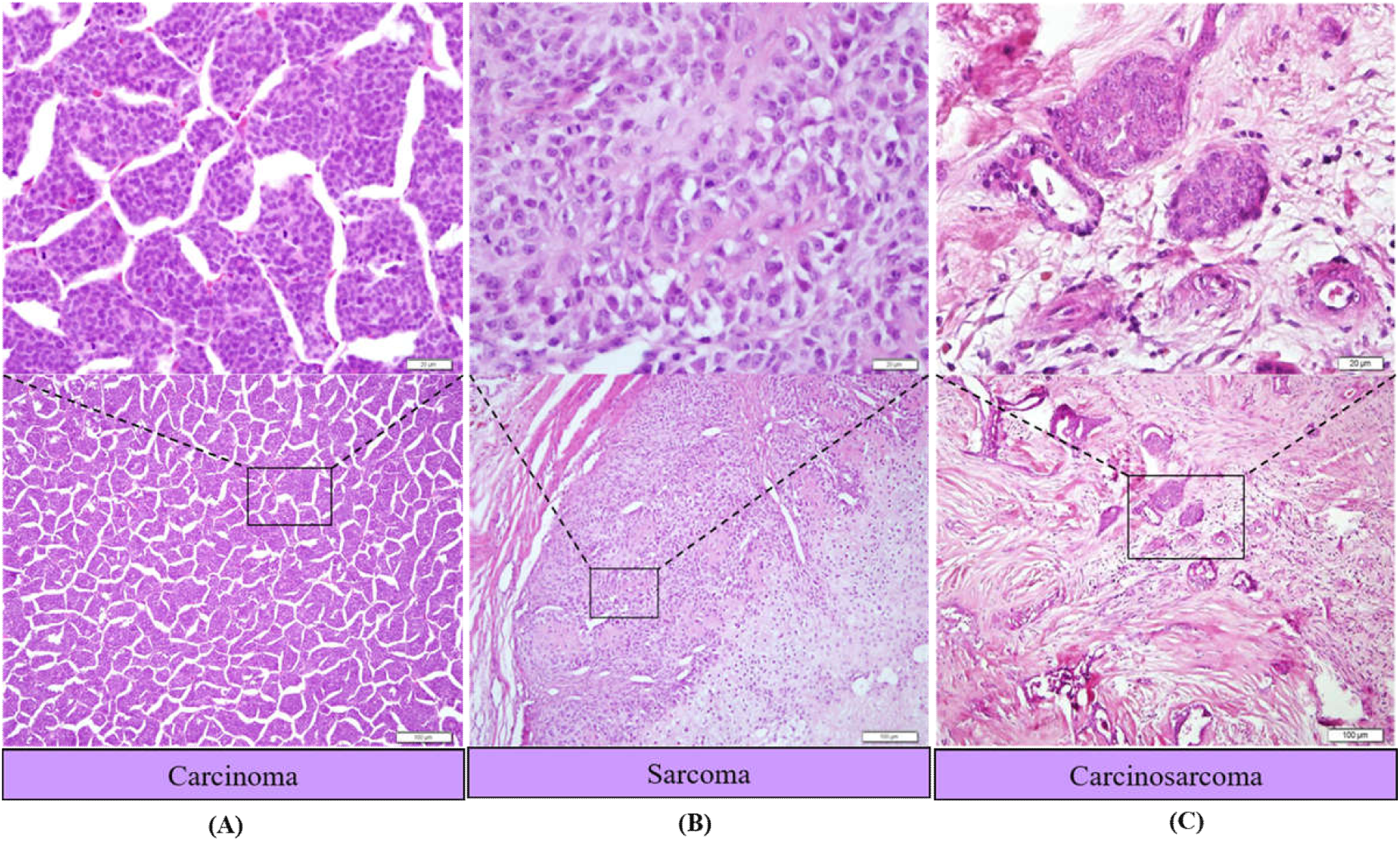
Canine mammary cancer A. Carcinoma, Low power lens: The gland epithelial cells are arranged in the island, High power lens: Glandular epithelial cells are arranged closely, with unclear boundaries, **B**. Sarcoma, Low power lens: Local areas showed cartilage-like, bone-like structure, most of them as an island. High power lens: The cells that in the outer edge of the cartilage-like, bone-like structure **C**. Carcinosarcoma, Low power lens:The tumor is not a complete capsule, composed of a large number of cell components, High power lens:Tumor cells varied. Glandular epithelial cells arranged in a group like, the cartilage-like structure can be seen. Scale bar = 100 and 20 μm. The top panel shows the enlarged view of the areas directed.

In addition to our analysis of somatic single nucleotide variants(SNVs), we also looked for indels. In total, we identified 16648 insertions and deletions (Indels) comprising 28 Indels in coding regions. The three most frequently mutated genes (indels mutations) *ACTB, CHD2*, and *EFL1* with mutations in the form of frameshifts indels are also prevalent in these genes **(Table S4)**. According to Gene Ontology (GO) analysis, these Indels are over-represented in 9 biological processes **(Table S4)**. Among these significant terms, most terms are highly associated with chromatin binding, DNA-templated, and cell differentiation.

### Characterization of somatic copy number alterations for canine mammary cancers

CNAs can affect gene dosage by altering the number of gene copies in the genome. Next, we analyzed the copy number profiles of canine mammary cancer. We identified recurrent large and whole-chromosome gains and losses in canine mammary cancer. We subsequently estimated the cellularity for the canine mammary cancer samples by using the widely-used software sequenza. The estimated cellularity range from (0.1-0.7) and the average sample cellularity is 37.2%, indicating the low purity of canine mammary cancer samples. We identified the chromosomal ploidy change for each chromosomal segment according to the read depth and B allele frequency revealed by whole-genome sequencing. The ploidy of the chromosomes in canine mammary neoplasms is generally diploid, with only three exceptions which have aneuploidy. Two solid carcinoma DB4-L4 (4), DB30-R3-R4 (3.2), and one osteosarcoma DB3L4-2 (3.7). It seems solid carcinoma can get aneuploidy. We also characterized somatic copy number alterations (SCNAs) compared to normal genomes, which is leading to 7762 chromosome segments ranging from 100bp to 1Mb including copy number gain (5889) and loss (1874) or LOH (loss of heterozygosity) affecting hundreds of genes per tumor. Five chromosomal CNAs are observed in three specimens on chromosomes 12, 16, 19, 22, and 29 **(Table S1) (Figure S2)**.

We identified a total of 9136 CNAs at a median of 338 CNAs per sample **(Figure S6)**, canine mammary cancer, and almost all the gains and deletions were observed in at least four samples **(Figure S2)**. GISTIC 2.0 analysis (with a threshold of q < 0.25) revealed 3 focal amplifications and 4 focal deletions recurrently altered in canine mammary cancer along with 25 recurrently altered whole arms **(Figure 2C and Table S6)**. Among the recurrent focal regions, 26.26, 27.27, 1.1, 3.3, 5.5, 8.8, 10.1,12.12, 16.16, 17.17, 18.18, 20.2 and 22.22 deletions have been previously found in canine mammary cancers. Deletion events occurring at 26.26 and 2.2 encodes an oncogene *PTEN* and *CDK7* which has been reported as a frequently amplified gene in canine mammary cancer^14^.

**Figure 2.**
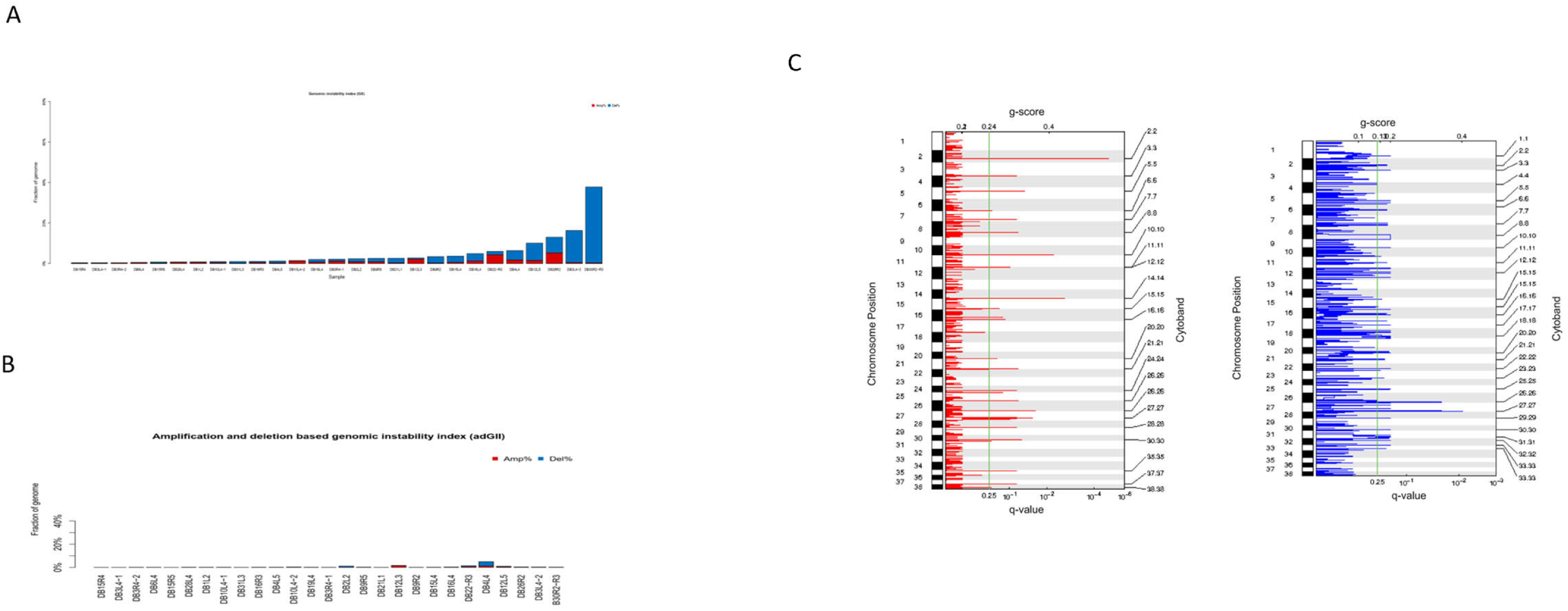
Genomic instability and copy number landscape of canine mammary cancer. A bar plot representing the fraction of the total genome altered with copy change ≥1 relative to median integer ploidy, which is termed as the genomic instability index (GII). B. A bar plot representing the fraction of the total genome affected by high-copy gains and losses (amplification and deletions with copy change ≥2 relative to ploidy), which is termed as the amplification- and deletion-based genomic instability index (adGII). **C**. Amplification (Left) and deletion (Right) plots using all data and amplitude threshold. The green line represents the significance threshold (q-value = 0.25).

To detect the potential functional effect of recurrently focal events, we performed Gene Ontology (GO) analysis for the genes located in the amplified and deleted regions **(Table S7)**, respectively. Among the significant terms of amplified genes **(Table S7)**, nucleoside triphosphate catabolic process, cell cycle, and apoptotic processes were highly associated with tumorigenesis. On the other hand, the deleted genes were enriched, cell cycle in some protein serine/threonine kinase activity, and lipid catabolic process **(Table S7)**. These results highlight the potential functional roles of recurrent CNAs in the formation of canine mammary cancer.

Amplified and deleted genes the were enriched in **(Table S7)** polynucleotide 5’-phosphatase activity, cell-cell adhesion, methylguanosine mRNA capping, transcription coactivator activity, and neuropilin binding **(Table S7)**. In this way, we identified several genes found in the Cancer Gene Census (CGC) catalog that were focally amplified or deleted in canine mammary cancer. Specifically deleted genes *PTEN, CDK7, SPP1, UBE4B*, and *ARG1* were enriched in the regulation of transcription. These results highlight the potential functional roles of recurrent CNAs in canine mammary cancer.

We also measured genomic instability index (GII, defined as the fraction of the genome altered by CNAs, copy change≥1 relative to ploidy), and observed that the majority of tumors showed low genomic instability (median of 2.1% per tumor, Figure 2A) except two samples display high genome instability index DB30-R3-R4 (39.9%) and DB3L4-2 (18.9%) **(Table S1)**. It depicts that the canine mammary cancer shows genome stability. A median of only 0.15 % of the genome was affected by high-copy gains and losses (copy change ≥2 relative to ploidy; defined as amplification and deletion-based genomic instability index (adGII); **Figure 2B**).

### The landscape of mutational signatures in canine mammary cancers

Mutational signatures are the particular patterns of mutations which is caused by mutagenesis processes such as infidelity of DNA replication, repair deficiency, enzymatic DNA modification, endogenous/exogenous mutagen exposures, provides insight into potential biological mechanisms of carcinogenesis, which has exhibited applicability in cancer etiology, prevention, and therapy. Mutational signatures in canine mammary cancer have not previously been formally investigated. Here, we performed mutational signature analyses in canine mammary cancer and compared them with human breast cancer.

The mutation spectrum in 6 base substitution catalogs revealed a significant dominance of C→T/G→A transitions as in most tumors in humans **(Figure 3A)**, indicating similar mutation mechanisms (e.g., deamination of 5-methylcytosine) during tumorigenesis in both species. By performing the non-negative matrix factorization (NMF) algorithm and model selection approach in MutationalPatterns package **(see Materials and methods)**, we reported four distinct mutational signatures extracted from 27 canine mammary cancer specimens, comprising 3 previously observed signatures in human breast cancer and a novel mutational profile uniquely occurred in dog mammary cancer **(Figure 3B)**.

**Figure 3.**
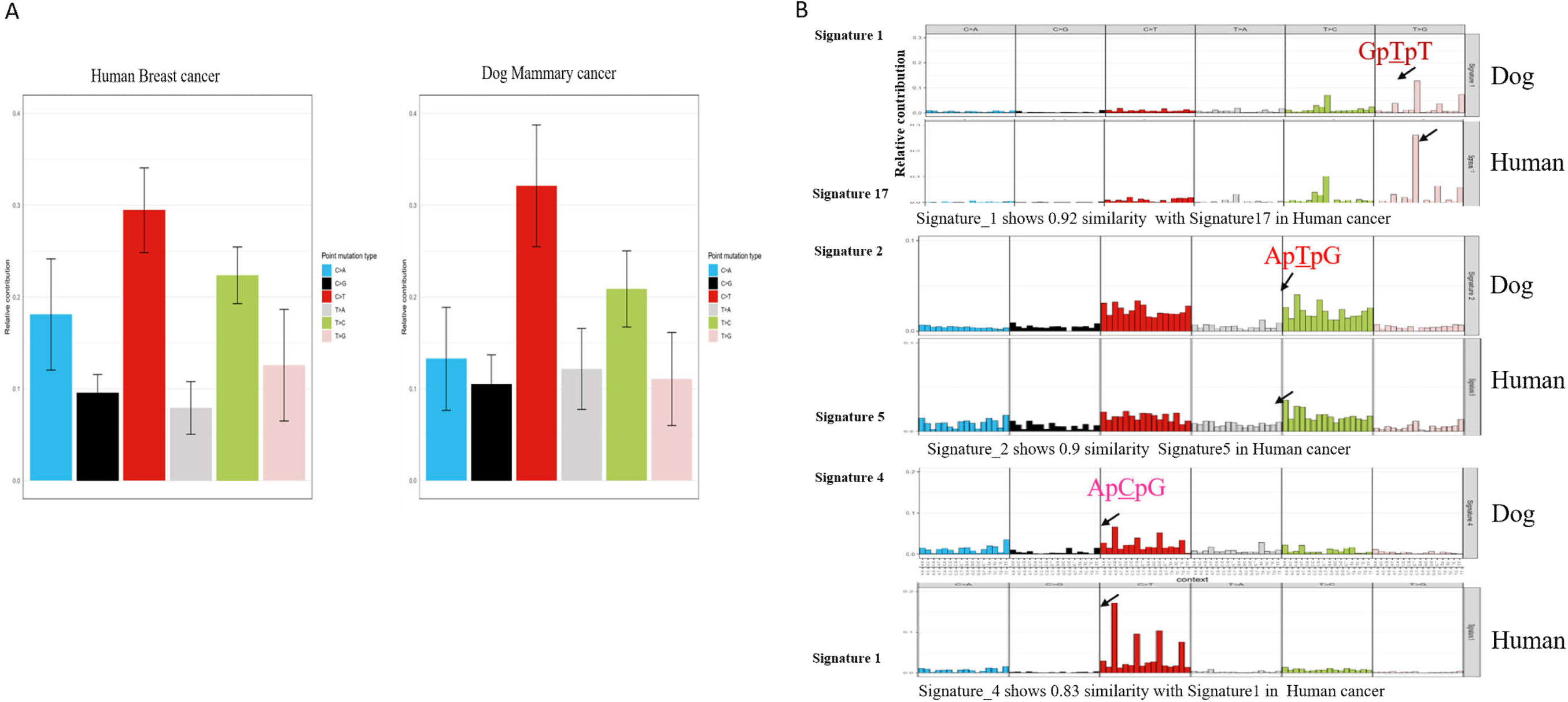
**A**. Mutation spectrum across human and canine mammary cancer. Mutation spectrum six base substitution types. Bars represent the mean relative contribution of each mutation type and error bars depict standard deviation. **B**. Comparison of mutational signatures in human and canine mammary cancer. Three mutational signatures in canine mammary cancer show similarity with human breast cancer (a): Signature 1 in dog shows 0.92 similarities with signature 17 in humans. (b): signature 2 dog shows 0.9 similarities with signature 5 in humans. (c): The signature 4 in dog shows 0.83 similarity with, signature 1in human.

Signatures 4 and 2 identified in dog cancer were analogous to human aging-related mutagenesis (COSMIC signatures 1 and 5), with cosine similarity of 0.83 and 0.9, respectively. Signature 4 features a predominance of C>T transition in the context of NpCpG trinucleotide, whose analogy in human displays etiology of spontaneous deamination of 5-methylcytosine, an epigenetic regulatory mechanism with implications for aging, and correlates with the age of cancer diagnosis in a clock-like manner ^15^ which may support the hypothesis of mutation accumulation theory ^16^, i.e., somatic mutational signatures during tumorigenesis have been acquired over the lifetime of individuals in both species. Based on the fact that the proportion of the enriched mutation types of NpCpG is much less in signature 4 in dog compared with signature 1 in humans (4% vs 11%), implicating a potential divergence in aging-related evolution during tumorigenesis. Signature 2 displays enrichment for T>C and C>T substitutions, closely resembled with signature 5, which has a predominance of in the ApTpN context with transcriptional strand bias.

Signature 1 in dog shows high similarity with Signature 17 with cosine similarity of 0.92, it is characterized by the prominence of T>G mutations at NTT trinucleotides and was previously identified in breast, stomach and oesophageal cancers but its aetiology is still unknown. Another pervasive mutational signature identified, Signature 3 is unique in canine mammary cancer which is primarily characterized by C>T has a high proportion of TpCpT trinucleotides substitution. Besides, C>G substitution at NpTpA mutations is enriched as well as T>A substitution in the TpTpA context shows the contribution to this novel mutational signature in canine mammary cancer. Signature 3 shows the contribution of almost all cancer samples. These results revealed that different mutational processes were operative during the progression of canine mammary cancer.

### Identification transcriptome signatures and DEGs in canine mammary cancers

RNA-seq analysis revealed total 1115 genes, of which 532 and 583 genes were up and down-regulated, differentially expressed (p≥0.01 and fold change log2fc > 2) between all types of canine carcinomas **(Supplementary Table9 and Fig. 4)**. Strikingly, among these genes, negative regulation of hormone secretion and tumor necrosis factor-activated receptor activity is the most significantly enriched functions **(Supplementary Table10)**. To understand the alteration in mammary cancer, we first investigated canine four normal mammary glands where both luminal and myoepithelial cells are visible. Additionally, 31 canine mammary cancer RNA sequences **(Table S8)** were used to identify the number of transcripts were listed in **Table S8**.

**Figure 4.**
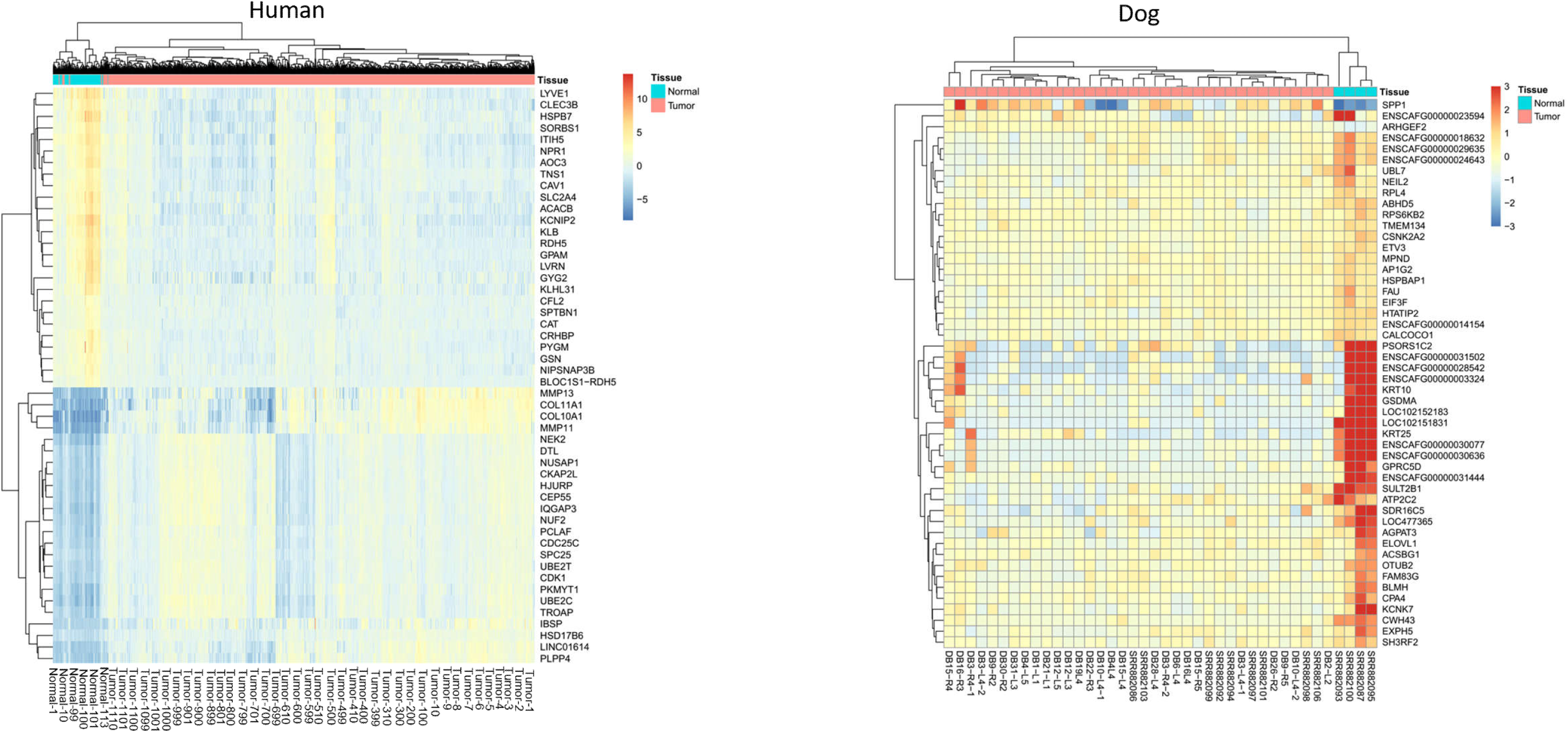
Cross-species comparison of the heatmap and hierarchical clustering of top 50 genes differentially expressed transcripts (p≥0.01, 2.0 FC) in human breast cancer and canine mammary cancer (red: upregulation; blue: downregulation) from the RNA-seq data. The left panel illustrates the human breast cancer and the right panel illustrates the canine mammary cancer. Each column represents a sample and each row represents a transcript. Expression level of each gene in a single sample is depicted according color key.

For the differentially expressed gene (DEG) analysis, we performed a comparison between thirty one including three subtypes of canine mammary cancer and matching adjacent four normal tissues. DEGs with a p-value < 0.01 and changes greater than 2-fold were determined for each comparison. Deseq2 analysis identified 1115, of which 532 and 583 genes were up- and down-regulated, respectively, in a comparison of the thirty-one canine mammary cancers and four matching adjacent normal tissues **(Table S9)**. Hierarchical clustering with the Kendall correlation matrix of the 1115 DEGs successfully distinguished canine mammary cancers and matching DEGs adjacent normal in a heat map analysis **(Figure 4)**. Our results indicated that transcriptomic signatures for canine mammary cancer subtypes (carcinoma, sarcoma and carcinosarcoma) might represent human breast cancer for few genes and provide new candidates for biomarkers. We did not observe a significant correlation among human breast cancer and canine mammary cancer **(Table S9 and S10)**. We found the different profiles of differentially expressed genes across the species.

Many cell cycle, cell adhesion, T-cell proliferation, and B-cell receptor signaling pathway pathways have been reported in canines **(Table S10)**. To better understand canine mammary cancer and human breast cancer, we performed KEGG pathway analysis using the web-based DAVID functional annotation tool (https://david.ncifcrf.gov/summary.jsp). For the pathway analysis, we used a list of DEGs genes from the overall canine mammary cancer comparison. Out of 2466 DEGs, 1587 up- and 879 down-regulated DEGs in human breast cancer were isolated and subjected to KEGG pathway analysis **(Table S10)**. Canine mammary cancer has been proposed as a comparative model for spontaneous tumors of human breast cancer. However, some new Ensembl genes were identified in canine mammary cancer and some of known cancer as a down-regulated in the dog but are known to be up-regulated in human breast cancer. These results are not consistent with previous studies, because we studied all subtypes of canine mammary cancer^17^. These results might represent similarities and discrepancies that exist between human breast cancer and canine mammary cancer.

### Comparison between human and canine mammary cancer

Following the analysis of canine mammary cancer, we proceeded to explore the catalog of mutated genes in each species. We turned to comparative genomics, taking advantage of the observation that a histologically similar malignant tumor, is common in dogs, which would enable discovery of actionable targets by sequencing of paired tumor-normal tissue samples. To compare somatic substitution of canine mammary cancer with human breast cancer, we profiled the mutational spectrum of SNVs. We analyzed 1080 WES from human breast cancer (TCGA data). Human breast cancer had in total 322,199 somatic mutations, consisting of 103,997 exonic mutations (68343 nonsynonymous, 24584 synonymous, and 5823 stop gain) **(Table S5)**. Intriguingly, the average number of coding region mutations per cancer case is approximately 9 in dogs, while 65 in human mammary cancer **(Figure 5A)**. After checking the average number of somatic mutations per tumor, we checked the mutation per Mb of the genome, which represents the accumulation of somatic mutations over the life of the tumor. Somatic mutation burden may influence tumor behavior and the mechanisms underlying the initiation and progression of cancer. The prevalence of somatic mutations was highly variable between and within cancer classes, ranging from about 0.001/Mb to more than 400/Mb. Breast in humans: average 1 mutation per megabase, ranging from 0.01⃜10. Intriguingly, the average number of coding region mutations per cancer case per megabase (Mb) is approximately 10 in dogs. It was notable that canine mammary cancer shows lower (2.67 and 0.187 mutations per megabase (Mb), respectively **(Figure 5B)** ^18^. Furthermore, we compared the genome-wide average mutation frequency per tumor per Mb in human and dog **(Figure 5C)**. These results revealed a low single nucleotide mutation burden in canine mammary cancer compare to human breast cancer which depicts that dog may need fewer mutations for triggering carcinogenesis ^19^.

**Figure 5.**
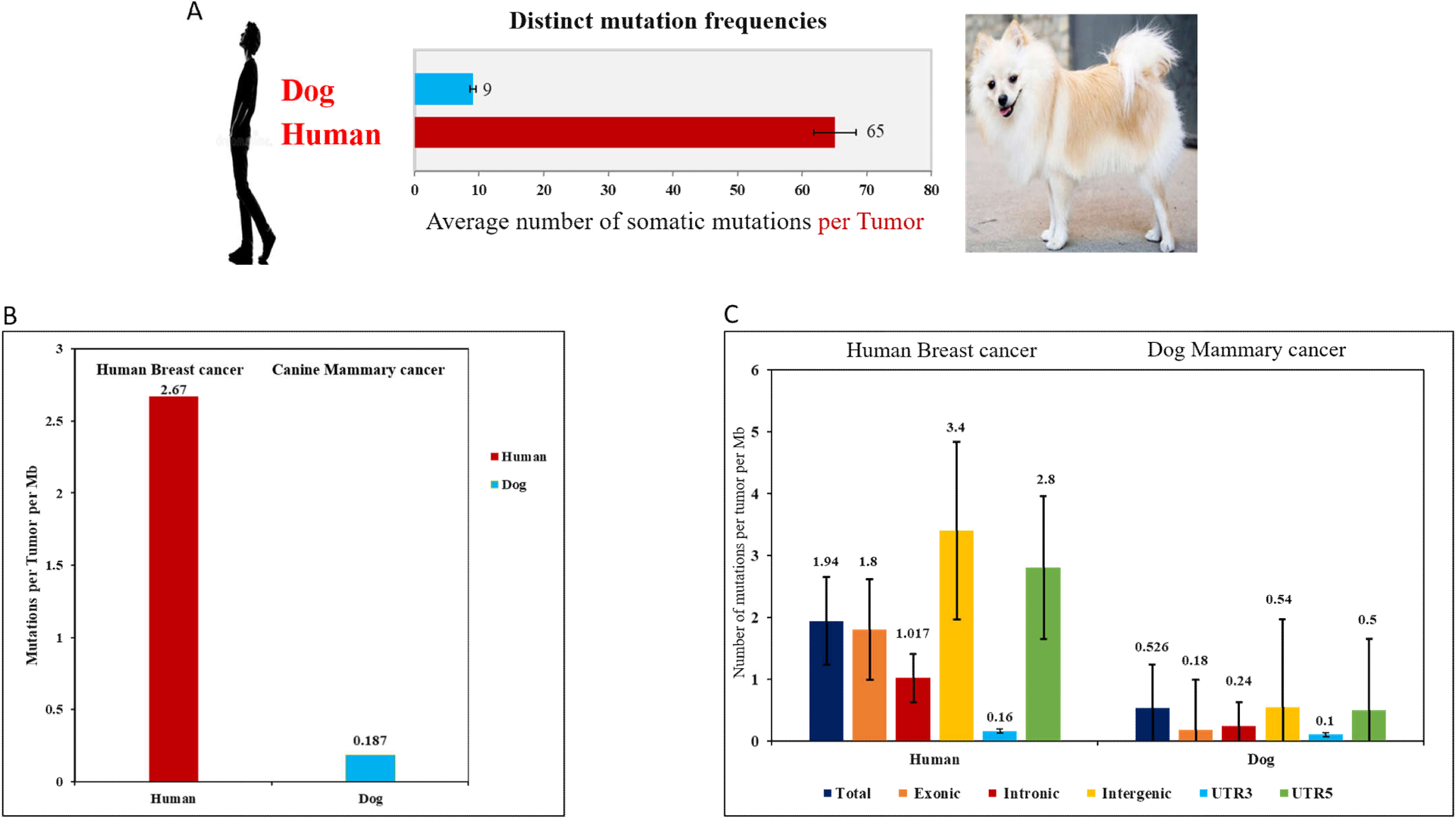
A. Average number of somatic mutations per tumor (Exonic region). Bars represent the average number of somatic mutations per tumor is 9 and 65 in the exonic region in human and canine mammary cancer, respectively. **B**. Cross-species comparison of average no. of mutations per tumor per Mb across the exonic region. Bars represent average no. of mutations per tumor per Mb across exonic regions in human and dog and error bars depict standard deviation. **C**. Cross-species comparison of genome-wide average no. of mutations per tumor per Mb Bars represent average no. of mutations per tumor per Mb across all classes of somatic mutation exonic, Intronic, intergenic UTR3, and UTR5 regions, and error bars depict standard deviation.

We have different DNA copy number profiles in dogs, it has fewer copy number changes compare to human breast cancer. However, by comparing chromosomal ploidy results, we found human breast cancer and canine mammary cancer show difference in ploidy. Chromosomal ploidy changes of the chromosomes estimated by sequenza in canine mammary neoplasms is nearly diploid on average, few tumors show the change in ploidy, significantly deviates from the findings of either aneuploidy or tetraploidy frequently observed in cancerous cells in human including breast cancer ^20^. Additionally, amplification and deletion were also observed over cancer genes previously implicated in breast cancer development including *PTEN, RB1, CDK6, AKT2, CCNE1, DNMT3A, MDM2, CCND1, ZNF217, ESR1, ZNF703, EGFR, IGF1R, AKT1, CDKN2, NF1*, and *CDH1* **(Figure S1)**.

### Similarity and divergence between human and canine mammary cancer genes

To compare cancer genes in humans and dogs, we explored the catalog of mutated genes in both species. A total of 179 genes from canine mammary cancer was compared with 1997 human breast cancer genes having mutation frequency >1%. The four most significantly mutated cancer genes are *PIK3CA* (32.2%) involve cancer initiation and progression, *TP53* (25.6%) regulate the cell cycle and *TTN* (25.6%) regulate ATP binding to the kinase in human breast cancer and *MUC16* (13.9%) plasma proteins protect from infectious agents **(Table S3)**, seems quite different from the dog mammary cancer as expected. The *PIK3CA* gene is one of the most frequently mutated common oncogene in human breast cancer (32.22%) and canine mammary cancer (33.33%). By performing Fisher’s exact test, we found three significantly different genes both in human and dog mammary cancer (Fisher’s exact test, P-value <= 0.028). *TP53* and *TTN* mutations have relatively high mutation frequency in human breast cancer compare to dog and *CKB* mutations predominate in canine mammary cancer. *NEB, MUC16*, and *PCLO* mutations were found to co-occur. Since we have less number of samples from dog mammary cancer compare to human breast cancer, we have limitation for this comparison. Thus, for the further comparison based on significantly mutated genes, we have chosen 127 SMGs (significantly mutated genes) from 20 cellular processes involved in cancer which already reported across 12 major cancer types in humans ^21^. These SMGs are involved in a wide range of cellular processes, including histone modifiers, genome integrity, receptor tyrosine kinase signaling, transcription factors/regulators, mitogen-activated protein kinases (*MAPK*) signaling, cell cycle, phosphatidylinositol-3-OH kinase (*PI3K-Akt*) signaling, histones, ubiquitin-mediated proteolysis, *Wnt/b-catenin* signaling, and splicing ^21^. Comparison of SMG in human breast cancer and canine mammary cancer **(Table 1)** shows the *PI3K-Akt* signaling pathway is significantly common in both species, which play an essential role in the initiation and development of mammary cancer and is considered to be a potential therapeutic target. We found many similarities, however, the genomes of canine mammary cancer lacked mutations in other key human breast cancer drivers, such as *TP53* and *CDH1*, etc. Thus, canine mammary cancer represents similarities and as well as a difference in mutation profiles from human breast cancer.

Interestingly *PIK3CA* frequently mutated the gene in both species, human and dog. We revealed a common occurrence of the same mutations in the *PIK3CA* gene, at position chr34, 12675674, which is identical to the *PIK3CA* mutation reported with high frequency in human tumors (8% of all human tumors). The mutation was present in 87.9% of canine mammary tumors and shows a protein truncating effect **(Figure 6)**. and shows the protein truncating effect, the amino acid substitutions at conserved sites **(Figure 6)**. The amino acid substitutions H1047R results in enzymatic overactivation^22^. This conservation for both amino acid substitutions H1047R and H1047L, strongly suggests that the same mutations in the canine cases also play a functional role in mammary cancer.

**Figure 6.**
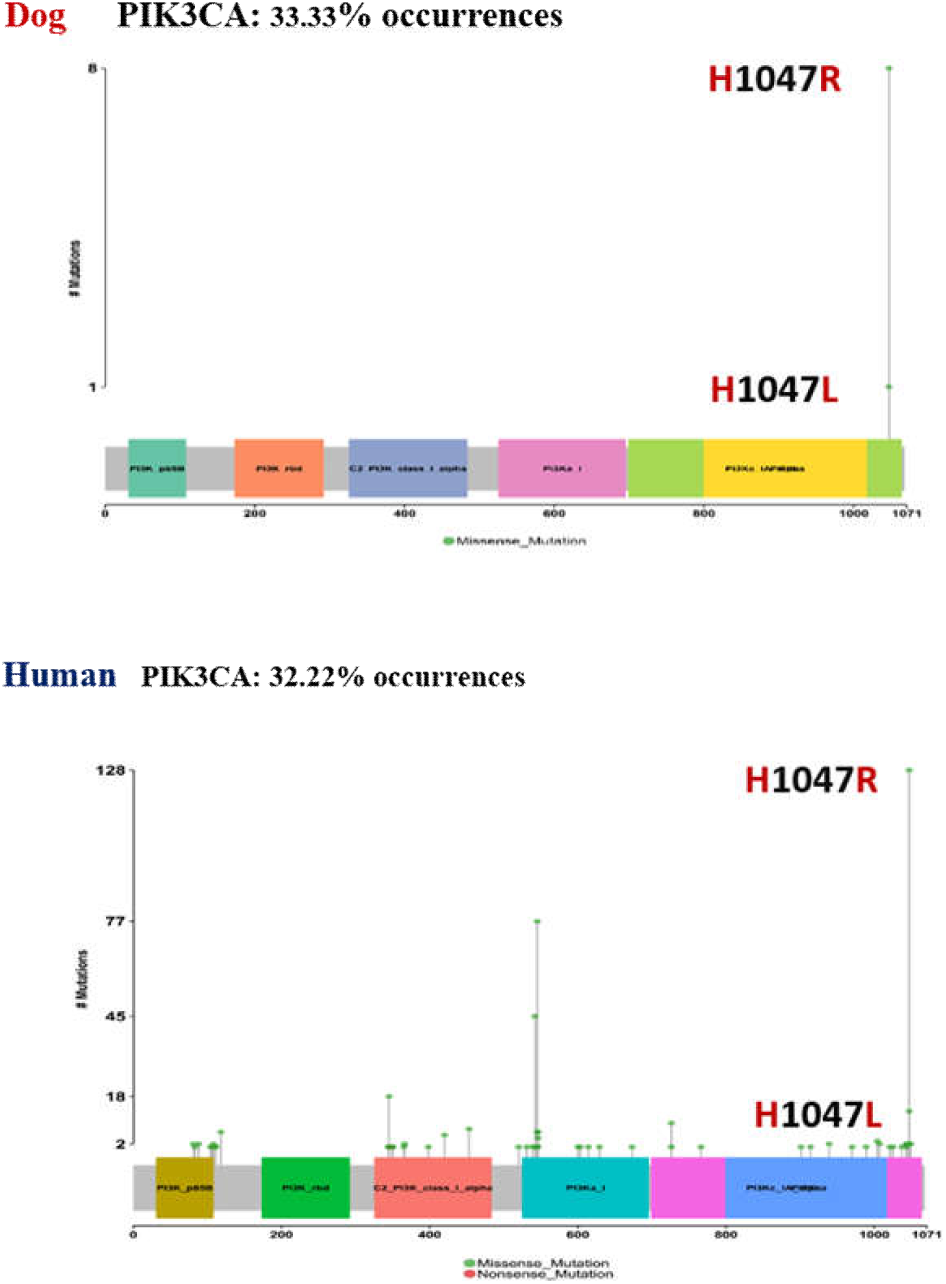
Alterations in the *PIK3CA* gene. The mutations needle plot shows the distribution of the observed cancer mutations along the protein sequence. The needles’ height and head size represent mutational recurrence. PIK3CA is frequently mutated and shows a protein-truncating effect in humans and dogs.

To investigate the selective pressure acting on the same tissue across human and canine mammary cancer, we choose distributions of the evolutionary rate (*Ka/Ks*) to study this phenomenon. The intensity of selection in cancer cases can be expressed as the *K*_*a*_/*K*_*s*_ ratio, where *K*_*a*_ is the number of nonsynonymous changes and *K*_*s*_ is the number of synonymous changes. We calculated the genomic *K*_*a*_/*K*_*s*_ ratio for both human and dog cancers and found that *K*_*a*_/*K*_*s*_ in human equals to 1.12, a hallmark of neutrality, indicating a generally neutral evolution. Whereas *K*_*a*_/*K*_*s*_ in a dog is equivalent to 0.67, implying a relatively strong negative selection in coding regions **(Figure 7)**. Most tumors have been evolving neutrally in humans, but here negative selection in dog indicates, maybe coding point mutations are lost through negative selection in canine mammary cancer or caused by the lineage-specific characteristics ^23,24^. Here, we found the evidence, which implies that different organisms have different selective pressure on the same organ.

**Figure 7.**
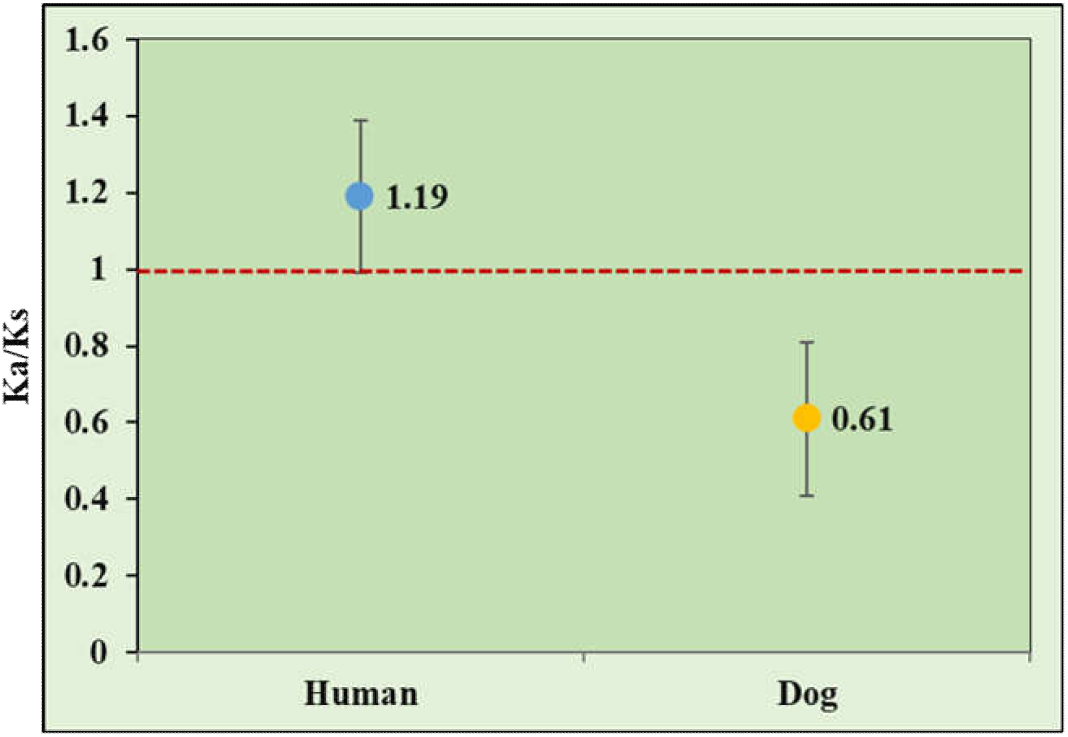
Genome-wide Ka/Ks value in human breast cancer and canine mammary cancer. Distributions of the genome-wide evolutionary rate (Ka/Ks) dog versus human mammary cancer. Y-axis represents the Ka/Ks value and the X-axis represents the organism name.

### Assumptions for varieties of mutations numbers between dog and human breast cancer The antagonistic pleiotropy theory

An evolutionary term (antagonistic pleiotropy theory) says benefits in early life increase the organism’s fitness, which may make it disease susceptible later in life. This explains the reason why the degree of cancer susceptibility varies across mammals. As, humans live longer than dogs, they have acquired the biological ways to slow down and regulate somatic evolution increase the cancer risk in humans. Human life expectancy worldwide has increased dramatically due to improved food, living conditions, hygiene, and medical care. The dog due to having a small life span cost of maintaining reproductive capability, and of reproduction itself, has been demonstrated in a wide range of animal species. However, little is understood about the mechanisms underlying this relationship. We have explored the aging genes showing selection in humans compared to dogs^16^. We found that sterilization was strongly associated with an increase in lifespan, and while it decreased the risk of death from some causes, such as infectious disease, it increased the risk of death from others, such as cancer. If the female dog is sterilized having less chance of mammary cancer, as we observed in our datasets only two female dogs are sterilized and having mammary cancer. Our dataset includes 11 dogs having age 12 or >12. In general, smaller breeds of dogs tend to live longer and may age slower. Dogs are considered old after they are about 8 years old and having chances to get cancer, hearing loss, cataracts, and senile dementia are also observed. To address this question, we compared 237 aging-related genes both human and dog the aging-related genes do not show selection in dog mammary cancer. As the previous analysis shows, Maximum lifespan potential (MLSP) estimations show that longevity did not evolve in dog lineage^25^. Natural selection acts on species when longevity evolves, give insights into adaptive genetic changes associated with the evolution of longevity in mammals^13^. It seems by experiencing artificial selection for breeding dog trade-off its defense mechanism DNA repair, immune function, and cell cycle genes. It is the evidence as certain mechanisms that are beneficial for dogs in breeding reproduction and growth, early in life could lead to bad health effects later similar to humans. Pleiotropic effects emphasizing the importance of life span when studying cancer across species. The dog has undergone human-directed breeding and artificial selection, which give rise to phenotypic variants many molecular mechanisms are involved in dog mammary cancer progression as it shows a variety of cells. In addition, Dog develops some horrible cancer that is absent or rare in humans for example transitional cell carcinoma of the bladder, histiocytic sarcoma, squamous cell carcinoma, and canine venereal transmissible tumors. The antagonistic pleiotropy theory shows that evolutionary adaptations maximizing fitness in dogs but increase disease burden after reproduction.

### Aging-related genes

To infer the antagonistic pleiotropy theory, we hypothesized that these genes can contribute to aging-related disease. To test this, we downloaded aging-related genes from http://genomics.senescence.info/download.html and calculated the Ka/Ks values for these mutated genes in both humans and dogs. For dogs, only 5 genes have nonsynonymous mutations, while for humans, the gene number increases up to 231. The 5 genes for dogs have Ka/Ks ranged from 0.64 to 1.43, where, only *PIK3CA* has Ka/Ks of 1.43 and the others have comparable Ka/Ks with the genomic (background) Ka/Ks (0.64-0.83). Whereas, the same set of genes in human represent Ka/Ks ranging from 1.64 to 12.96, far higher than the background Ka/Ks (1.16). Additionally, the Ka/Ks distributions for these aging-related genes are dramatically different between humans and dogs. Majority of the genes (221/237) didn’t experience positive selection (maybe experience negative selection) in the dog with all of the genes having Ka/Ks =0 or <1 with exception of *PIK3CA*, whereas, inhuman, 3 genes have Ka/Ks >10; 30 genes have Ka/Ks of 3-10; 32 genes: 2-3; 50 genes: 1-2; 43: <0.5, which indicated a positive and negative selection in human and dog, respectively **(Figure 8A)**. The *PTEN* is considered a gatekeeper tumor suppressor gene involved in multiple mechanisms that lead to cellular defense against neoplastic transformation in human breast cancer. It shows the highest Ka/Ks 12.96 as its occurrence is 3.3% in Human breast cancer **(Figure 8B)**. *PIK3CA* converge genetically between human and dog mammary cancer as its occurrence is high in both species. But, in humans it experiences selective pressures Ka/Ks is 2.37 but in Dog Ka/Ks is 1.43 seems lower as compared to humans, maybe neutral. So, it shows convergence and a lack of overall selection^23^. As, in all cancer cases many genes experience strong selective pressures but do not converge genetically. Alternatively, different cancer cases may be driven by weak selection, which may nevertheless converge. Convergence and selective intensity are distinct phenomena^1^. These findings supported by, antagonistic pleiotropy theory hypothesis, an evolutionary trade-off that evolutionary adaptations benefiting the fitness in early life increase disease burden in later life after reproduction. Similarly, we have found common genes in the same tissues in different species showing different selection pressure. Ecological factors of the mammary glands across species show, same organs in different mammals impose different selective pressures on the same set of genes^26^.

**Figure 8.**
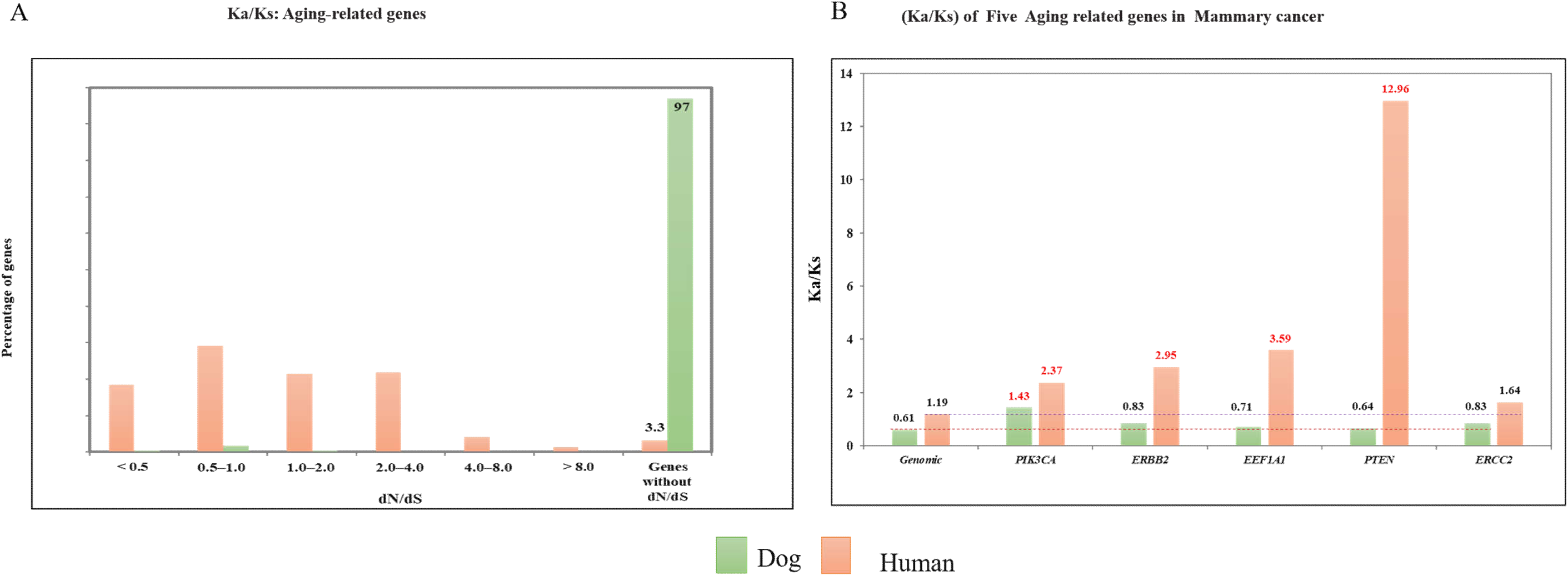
**A**. Ka/Ks of aging-related genes in mammary cancer Distributions of the evolutionary rate (Ka/Ks) of aging-related genes in mammary cancer (human versus dog). Genes without Ka/Ks (the genes are not mutated or have the only synonymous mutation) are 97% and 3.3% in dogs and humans, respectively. The observations show much higher percentages of genes with Ka/Ks in human breast cancer. **B**. Ka/Ks for Genome-wide and five common aging-related genes in mammary cancer Although the genome-wide average of Ka/Ks is very close to 1, the Ka/Ks ratio for individual genes deviates from 1 in human breast cancer. The genome-wide and the Ka/Ks ratio for individual genes is less than one in canine mammary cancer except for *PIK3CA*, which has Ka/Ks=1.43.

As these aging-related genes involved in development in the early life of a human, the mutation in these genes shows the selection in human breast cancer after reproduction. The modern human populations have substantially increased lifespan over the last two centuries and limited purifying selection, which has been weakened by the use of advanced medications. However, in the dog, it sounds different, as trade-off did not observe.

Dog life span did not increase relative to human and the genes involve in development, did not show selection in canine mammary cancer. Dog show different phenomena compare to humans may be in dog’s domestication trade-off the aging or cancer-related phenotypes^27^. Dogs have undergone thousands of years of human-directed breeding and contribute to cancer risk rather than aging selection. The present study represents the first report of testing the antagonistic pleiotropy theory hypothesis. Although most wild animals, including dogs, develop cancer, the rates are typically much lower than in domestic animals. Trade-off is important and helpful to study the origin of aging-related diseases^12^.

### Inter and Intra-heterogeneity of canine mammary cancers

Little is known about tumor heterogeneity when it comes to canine mammary cancers. To examine the functionally essential variations related to canine mammary cancer heterogeneity, we performed a phylogenetic analysis of six cases of multi-tumor mammary carcinomas by using single nucleotide variations (SNVs). The morphology and biological behavior of canine mammary cancer are quite heterogeneous. This is 1^st^ time we are going to report inter and intra-heterogeneity of canine mammary cancers.

Phylogenetic relationships based on SNVs accurately reflected the clonal origin, like our previous studies about human cancers shows that genetic variations can be used to reconstruct clonal evolutionary relationships among different tumors ^2,28,29^. Here, we found that inter and intratumor heterogeneity in canine mammary cancer is very high. Each canine patient has a different tumor evolution pattern, for example independently evolving, having a common origin, or genetically identical or unique tumors. Since. high level of heterogeneity within and between patients or tumors is a major obstacle to successful cancer therapy^30^. The histological and molecular heterogeneity, together with evolutionary patterns, demonstrate distinct biological behavior of these tumors, different clinical and therapeutic approaches are required for treatment^8^. In addition, dogs can help us to study inter and intra tumor heterogeneity, as each female has eight to ten mammary glands. It is rare for women to develop more than one tumor in the breast. In this study, we carry out whole-genome sequencing on six cases of multi-tumor mammary carcinomas. Two tumors exhibited the intra-tumor heterogeneity with multiple morphological patterns. Multiple tumors are evolutionarily more complex than single tumors as they have divergent microenvironments and experience cell migration. The cases of canine mammary cancer used in this study are summarized in **Table S12**. Further analysis revealed, four cases are of independently originated because tumors from the same canine patient share very few somatic mutations, for identical tumors >60% of mutations are shared. Independently originated tumors, are interesting, as they offer no information on the evolution of inter-tumor diversity within one canine patient^31^.

### Phylogenetic relationship of DB3

Dog3 showed two tumors, one on the left side fourth position (L4) and second on the right side the same position (R4) with mixed morphology. Histopathology results display DB3-R4-1 as carcinosarcoma, DB3-R4-2 as osteosarcoma, DB3L4-1 as lipid carcinoma and DB3L4-2 is carcinosarcoma. We have sequenced four samples from Dog3 show only one common mutation seems to evolve independently **(Figure 9A)**, showing a different pattern from the human. It demonstrates the heterogeneity of canine mammary cancer. The phylogenetic tree of Dog3 is interesting, it displays the highly heterogeneous nature and independent origin of these tumors within one canine patient. As, both tumors DB3L4 and DBR4 shows the carcinosarcoma, which contains two types of cancer cells (epithelial and mesenchymal). Thus, this phenotypic diversity in one-tumor shows may be epithelial to mesenchymal transition (EMT) that happens within the tumor in canine mammary cancer.

**Figure 9.**
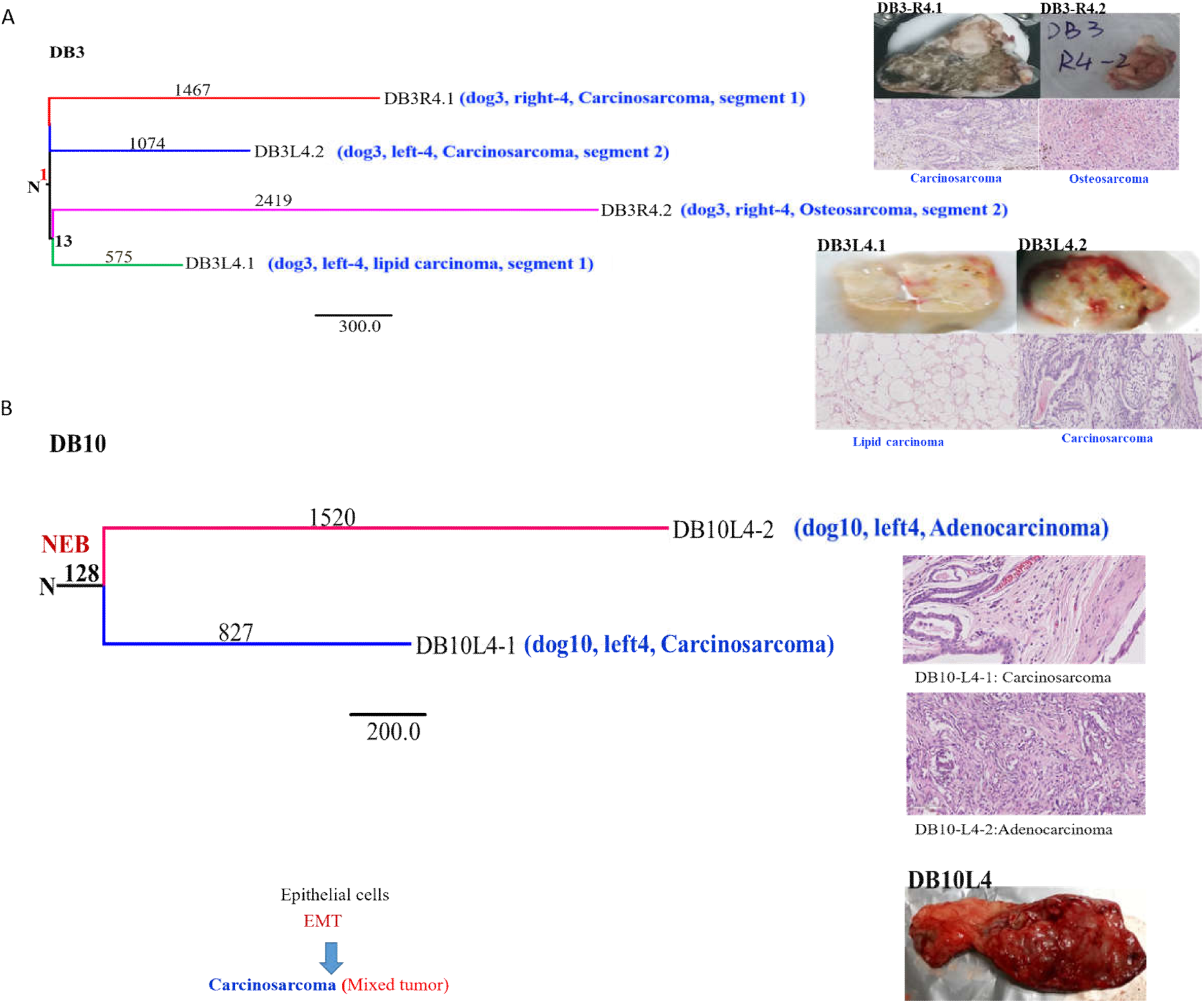
**A. Phylogenetic relationship between the tumors in Dog 3 between four samples.** The lengths of branches are proportional to mutation numbers. The tree was anchored using a DNA sequence from the normal tissue of a canine patient. **B. Phylogenetic relationship between two samples in Dog10**. The tree was anchored using a DNA sequence from the normal tissue of the canine patient. The lengths of branches are proportional to mutation numbers. The numbers correspond to the number of common mutations and specific mutations in the samples.

### Phylogenetic relationship of DB10

Dog 10 had one tumor one on the left side fourth position (L4) with diverse morphology (Figure 9B). Histopathology results show DB10L4-1 as carcinosarcoma and DB10L4-2 is adenocarcinoma. We have sequenced two samples from Dog10 tumor, shows 128 common SNVs (*NEB* gene shows exonic) mutations seem identical but the morphology is different. It seems epigenetics may also involve in this mixed morphology. Similar to dog3, may be epithelial to mesenchymal transition (EMT) happened within the tumor. Additionally, epigenetics changes may be contributing to the phenotypic diversity within the tumor rather than SNVs. We can assume from EMT the osteosarcoma originates, as shown in **(Figure 9B)** ^32^. By this process, we can also say maybe the first tumor may be the cause of the second tumor within the same canine patient. Despite the intratumor heterogeneity, we study the inter mammary tumor heterogeneity within one canine patient, as a female dog has eight to ten mammary glands. Six canine patients have multiple tumors. However, DB3 shows inter and intratumor heterogeneity, as we have explained above. The other remaining five canine patients with multiple tumors are discussed here.

### Phylogenetic relationship of DB4 and DB9

Dog 4 had two tumors on the left side, one on the left side fourth position (L4) and second on the left side fifth position (L5). This canine patient has two tumors with different histopathology and shared genome-wide 25 SNVs **(Figure S5A)**. It shows different histopathology does not share the common mutations. These results show, genetically unique tumors are evolving separately within one canine patient. Dog 9 had two tumors on the right side, one on the right side fourth position (R2) and second on right side fifth position (R5). Analysis of this canine patient shows similar histopathology for both tumors, a subtype of carcinoma, and shared genome-wide 98 SNVs **(Figure S5B)**. It demonstrates that carcinoma share genome-wide similar mutations. This canine patient shows both tumors are may be identical.

### Phylogenetic relationship of DB12 and DB16

Dog 12 had two tumors on the left side, one on left side third position (L3) and second on left side fifth position (L5). Both tumors are a type of sarcoma and a shared genome-wide 15 SNVs **(Figure S6A)**. It demonstrates that sarcoma may be shared genome-wide fewer mutations. These results indicate, genetically unique tumors are evolving separately within one canine patient. Dog 16 had two tumors same histopathology, one left side fourth position (L4), and the second tumor on the right-side third position (R3). Two tumors shared a few common genome-wide SNVs (16) **(Figure S5B)**. It shows sarcomas also share less genome-wide mutations similar to dog12. These results show tumors are evolving separately, therefore genetically unique in one individual.

### Phylogenetic relationship of DB15

Dog 15 had three tumors, one on the left side fourth position (L4)), second on right side fourth position (R4), and third on right side fifth position (R5). This canine patient has three tumors with different histopathology two tumors are the subtype of carcinoma (solid and simple carcinoma) and one mixed tumor. Three tumors shared genome-wide 9 SNVs. DB15 share the exonic mutation *PIK3CA* gene, which is a known driver gene, appear these tumors evolve from a common ancestor cell **(Figure 10)**. This result shows that mixed tumor which has epithelial and bone cells may be originated from epithelial cells. Additionally, maybe epithelial to mesenchymal transition (EMT) is cause diverse phenotypes in dog mammary cancer similar to dog3, dog4, and dog10. Canine carcinosarcoma can be the best model to study epithelial to mesenchymal transition (EMT), it can be assumed due to this process these tumors have different morphology. As, it is a natural behavior in canine mammary cancer, it also happens in human breast cancer but very rare. However, further studies are needed to explore the heterogeneity in canine mammary cancer by using multiple tumor regions sampling and multi-omics analysis.

**Figure 10.**
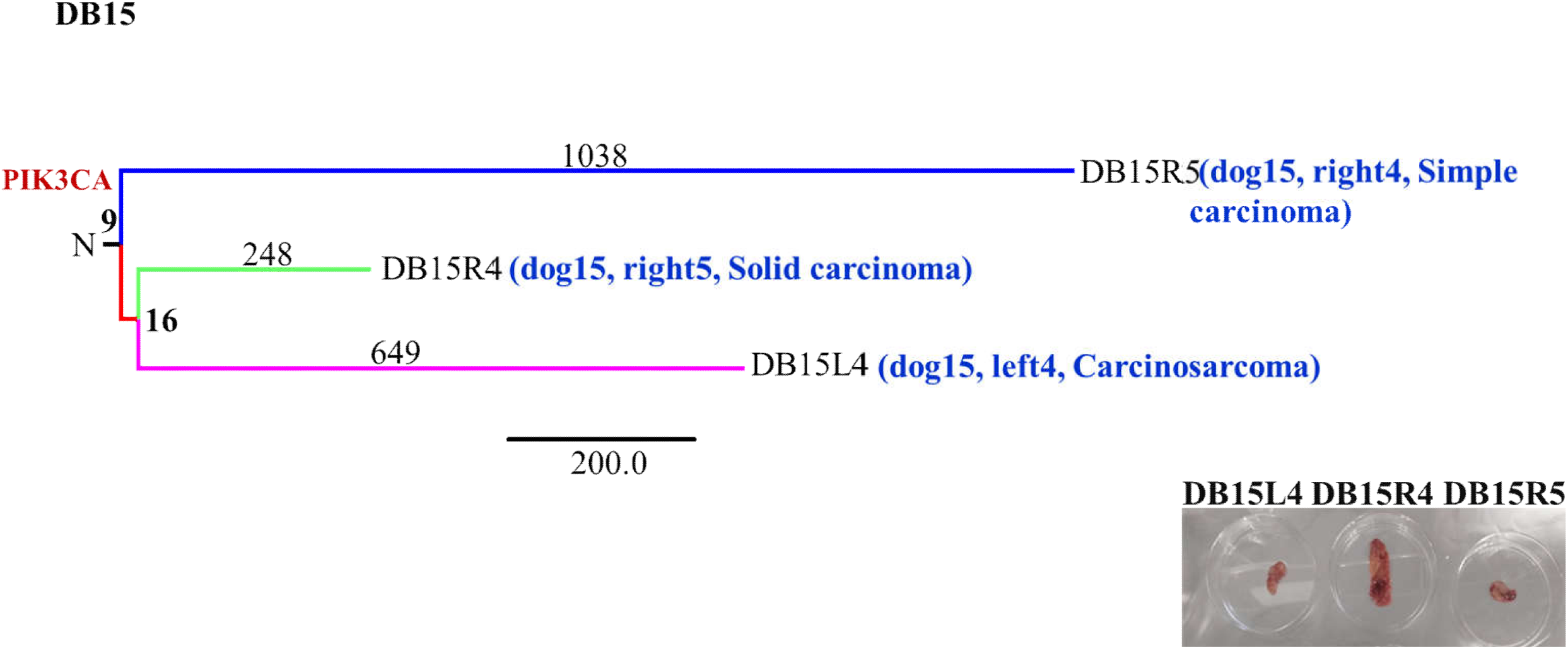
Phylogenetic trees based on SNVs of Dog15. The phylogenetic tree has been constructed using somatic mutations from three tumors (simple carcinoma, solid carcinoma, and carcinosarcoma). The lengths of branches are proportional to mutation numbers.

## DISCUSSION

The recent development of sequencing technology has opened the door for detailed profiling of canine mammary cancer genomes. Our study represents the first, as far as we are aware, to examine somatic genomic alterations in all subtypes of canine mammary cancer. We have performed a cross-species cancer genomics analysis, by using whole-genome and RNA sequencing a cohort of 27 cases from canine mammary cancer. We identified recurrent mutations in more than half of the cases whose human homologs are well-established cancer drivers, strongly suggesting that these candidates indeed drive canine. Taken together, canine and human studies highlight the benefits of comparative diseases genetics^5^, the importance of selection for genome evolution, as well as the extreme complexity of genome function and regulation. Nevertheless, the lack of a deep understanding of the genetic causes of tumor variation in canine cancer limits the use of this naturally occurring model for understanding human disease. Cancer in dogs also follows the general epidemiology of human cancer occurrence^7^. The prevalence of different cancer types between human and dog has provided some hints, the cross-species cancer genomics may offer new insights into cancer evolution. Through 100X WGS, we have, for the first time, characterized the genomic landscape of canine mammary cancer.

We identified and compared the mutation patterns of both human and dog, in terms of mutated genes and the number of mutations that may influence tumor behavior and the mechanisms underlying the initiation and progression of cancer. The average number of coding region mutations per case of cancer is higher in humans than in dogs. Dogs may need fewer mutations for triggering carcinogenesis ^33^. As dogs have been undergone strong artificial selection within the last hundreds of years with extremely high levels of morphological specificity based on strict breeding programs, leading to a considerable genetic disease such as cancer as mutations accumulated within the closed gene pool ^7^. It depicts the side effect of domestication has been the inadvertent enrichment for different types of diseases including cancer. Further, we reveal divergent indels and DNA copy number profiles, as compare to human breast cancer, canine mammary cancer showing fewer indels and copy number changes.

Importantly, we identified four mutational signature in canine mammary cancer. Three mutational signatures show the similarity with human breast cancer signatures. We found signatures 4 and 2 in canine mammary cancer were analogous to human aging-related mutagenesis (COSMIC signatures 1 and 5), respectively. Signature 4 analogy in human displays etiology of spontaneous deamination of 5-methylcytosine, an epigenetic regulatory mechanism with implications for aging, and correlates with the age of cancer diagnosis in a clock-like manner ^15^, which supports the hypothesis of mutation accumulation theory, i.e., somatic mutational signatures during tumorigenesis have been acquired over the lifetime of individuals in both species. Based on the fact that the proportion of the enriched mutation types is much less in signature 4 in dog compared with signature 1 in humans (4% vs 11%), implicating a potential divergence in aging-related evolution during tumorigenesis ^34^. Additionally, we found a novel somatic mutational signature in canine mammary cancer, previously not described in human cancers, these mutational signatures designate that dog share same mutagenesis processes like the human, however still different at the mutational level that might be the result of the difference in genetic background.

We identified the RNA expression, which suggests canine mammary cancer counterparts of human breast cancer, can provide new clues for biomarker screening for human breast cancer. However, some new Ensembl genes were identified in canine mammary cancer and some of known cancer genes as a down-regulated in the dog but are known to be up-regulated in human breast cancer. As, previous studies describe the canine carcinoma shows high similarity with human breast cancer but sarcoma and carcinosarcoma not. We thus determined transcriptome level across the specie not compatible at all. Since, many studies have been performed in comparison to human breast cancers and canine mammary at the transcriptome level. Thus, our results reveal the similarities and discrepancies between human breast cancer and canine mammary cancer^17^.

Further, we reveal patterns of selection in cancer evolution by adapting methods from evolutionary genomics and applying them to cancer genomes, which is characterized by the overall representation close to neutrality in human breast cancer^23,35^. Genome-wide Ka/Ks ratio for human’s equals 1.19, a hallmark of neutrality, indicating a generally neutral evolution and in the dog, Ka/Ks is 0.61, seems negative selection. As, most of the tumors in human have been evolving neutrally, which does not mean the absence of selection^23^. It only reflects that the effect of positive selection in accelerating evolution is exactly canceled out by the effect of negative selection ^6,23^. Moreover, negative selection in canine mammary cancer indicates, maybe coding point mutations are lost through negative selection or there are other unknown reasons. Purifying selection in the evolution within tumors. Natural selection can be positive (for the beneficial mutations) or negative/purifying (against the deleterious ones). Little attention has been paid to the power of negative selection in cancer evolution^36^. Wu et al. (2016) reported the average Ka/Ks ratio of 1 on all genes and the exceeding percentage of negative sites in comparison with the simulated distribution, suggesting the possibility of negative selection operating on the genes with the essential function of cells should not be ignored^26^. The operation results in stabilizing selection through the purging of deleterious variations that arise, eventually reducing the variation in phenotypes in populations. In the case of cancer cell population without recombination, the effect of negative selection could be high, as the purging of deleterious variants will result in the removal of linked variation. The test on the evolutionary forces and exploration of the effect and genes under selection remains required.

Further, we compared aging-related genes between human breast cancer and canine mammary cancer. Our results demonstrate that aging genes with somatic substitutions are showing a clear signal of positive selection, Ka/Ks is significantly high in human breast cancer as compared to the dog. Here, we get the answer to our main question, whether the same organs in different mammals impose different selective pressures on the same set of genes? These findings provide the answer to our question, the same set of genes in the same organs in different mammals impose different selective pressures^6^. Evolutionary constraints are a major theoretical challenge for explaining the linkage of genes in aging and cancer ^16^. Importantly, our findings with aging-related genes in humans can be explained by widespread antagonistic pleiotropy hypothesis which suggests that the same gene can be favorable for fitness but can confer the risk of traits later in life ^13,37^. This theory has not been rigorously tested previously.

But, in the dog, it looks different, as we did not observe selection on aging-related genes in canine mammary cancer. Therefore, maybe in dog’s domestication trade-off the aging or cancer-related phenotypes of modern dog populations than at any time of macroevolution. The modern human populations have substantially increased lifespan over the last two centuries ^38^ and limited purifying selection, which has been weakened by the use of advanced medications ^39^. A side effect of domestication has been the inadvertent enrichment for different types of diseases. This interplay between desired and deleterious traits, some of which are pleiotropic, has been particularly well used in the study of both canine and feline genomics. As a consequence, ∼50% of pet dogs develop cancer in their lifetime. Presumably, as a consequence, advancing age is the major risk factor for diverse types of loss of function, and highly prevalent chronic and killer diseases. Nevertheless, important questions remain to be addressed.

So, maybe in dog’s domestication trade-off the aging or cancer-related phenotypes of modern dog populations than in any time of macroevolution. Dogs have undergone thousands of years of human-directed breeding and selection, giving rise to the hypothesis that selection for desired phenotypes may have led to piggybacking of deleterious alleles that contribute to cancer risk. Most modern dog breeds only existed for 300 years and many are derived from small numbers of founders. We can say by experiencing the same environment parallel evolution is most apparent in genes for digestion and metabolism, neurological process, and cancer ^7,40^.

Further, we found that inter and intratumor heterogeneity in canine mammary cancer is very high. Each canine patient has a different tumor evolution pattern, for example independently evolving, having a common origin, or genetically identical or unique tumors. Dog9 shows genetically identical tumors within one canine patient, other dogs show genetically unique tumors within one canine patient. Four canine carcinosarcoma, maybe indicate epithelial to mesenchymal transition (EMT), it can be assumed due to this process these tumors have different morphology^32^. The histological and molecular heterogeneity, together with evolutionary patterns, demonstrate distinct biological behavior of these tumors, different clinical and therapeutic approaches are required for treatment. To diversity and heterogeneity of tumor cell types and states (such as metastasis and drug resistance) in both species, one or multiple tumors from a single patient are of necessity. Thus, further exploration and analysis of whole-genome sequences from dog mammary cancer will be required to complete our understanding of the somatic mutational basis of the disease. Furthermore, multiple region sampling can help us to investigate the heterogeneity and phenotypic diversity in canine mammary cancer. Additionally, there is still a need to explore the epigenetic or regulatory cause of phenotypic diversity.

## MATERIALS AND METHODS

### Sample collection

Twenty-four malignant tumor masses and adjacent non-tumor tissues were surgically resected by simple or regional mastectomy from 17 female dogs with diagnosed mammary gland neoplasms in the Veterinary Teaching Hospital of China Agriculture University and the Beijing Guanshang Animal Hospital. Samples were collected under the guidelines of animal use protocols and informed consent signed by dog owners. The 17 female dogs covered 8 breeds consisting of the Silver fox, Teddy, Cocker, Poodle, Dachshund, Pomeranian, Shih Tzu, and Mixed, with a sterilization rate of 12% and an average age of diagnosis of 12 (6-15) years old, 4 of which presented the clinical or radiological evidence of spread to nearby lymph node or distant metastases in lungs. The 24 tumors varied in sizes which ranged from 1.2 cm of diameter to 9.2 cm. According to the clinical TNM staging system of canine mammary gland neoplasms (Table S2) that was developed by the World Health Organization (WHO) and modified by Withrow and MacEwen (1996) ^41^, the tumors were staged as I-V based on tumor size and metastasis status. The detailed information on the dog breed, age of diagnosis, and tumor features such as anatomical sites, tumor sizes, metastasis status, and TNM staging were summarized in **Table S1**.

For each specimen of mammary gland neoplasm, two sections were excised for histopathological examination and Immunohistochemical (IHC) test of hormone receptors and growth factors, respectively. The rest part of the specimen was stored at −80°C, until being processed for DNA extraction. This study was approved by the Animal Research Ethics Committee of Beijing Institute of Genomics, Chinese Academy of Science.

### Histopathological examination and grading

Histopathological examination was performed by veterinary pathologists in the Veterinary Teaching Hospital of China Agriculture University. A representative section of tumor specimen was fixed in 10% formaldehyde, dehydrated in ethanol, and Xylene, embedded in paraffin wax, sectioned with 4 μm thick and stained with haematoxylin and eosin (Beyotime, Shanghai, China). Histological classification of the specimen was determined according to the WHO criteria for canine mammary cancer ^42^. According to the Elston and Ellis method ^41^, histological grading of the specimen was assessed as well-differentiated (grade I), moderately differentiated (grade II), as well as poorly differentiated (grade III). The histopathologic classifications and grading were described in Table S1.

### Immunohistochemical (IHC) analysis

To determine the presence of hormone receptors consisting of ER and PR, and the epidermal growth factor receptor 2 (HER2) in a 5μm-thick, formaldehyde-fixed, paraffin-embedded tumor section, IHC experiment was performed by following the standard protocols ^43^. The tumor section was incubated with primary antibodies, including Anti-ERp57 antibody (Abcam, ab13506) for ER, Anti-PR antibody - C-terminal (Abcam, ab191138) for PR, and human epidermal growth factor receptor 2 (HER2) Anti-ErbB2 antibody (Abcam, ab16901) for HER2 were obtained from Abcam (Cambridge, MA, USA). Images were taken with a CX31 optical microscope (Olympus, Japan). Immunohistochemical (IHC) analysis was performed by veterinary pathologists in the Veterinary Teaching Hospital of China Agriculture University. According to the hormone receptor status and HER2 status, we have characterized the canine mammary tumors into four subtypes: Luminal A (ER+, PR+, and HER2-), Luminal B (ER+, PR+, and HER2+), HER2-overexpressed (ER-, PR-, and HER2+), and Triple-negative (ER-, PR-, and HER2-)^44^ Table S1.

### DNA extraction, library construction, and WGS

Genomic DNA was extracted from the dissected cancerous or adjacent normal tissues using the QIAamp® DNA Mini Kit (QIAGEN, Duesseldorf, Germany). DNA quality and concentration were monitored using agarose gel electrophoresis, Bioanalyzer 2100 system (Agilent Technologies, CA, USA), and Qubit® 2.0 Fluorometer (Life Technologies, CA, USA). High-quality sequencing libraries were constructed by following the protocol of IlluminaTruSeq™ (cat. no. FC-121-2003) DNA preparation kit (Illumina, CA, USA), then were sequenced on Hiseq2000 platform (Novogene, Beijing, China). Approximately 114∼150 millions of clean paired-end reads with a length of 150 bp were generated per sample. The average sequence coverage was 90-fold for both tumor and normal samples (Table S1B). Genome sequence data has been deposited in the genome sequence archive of Beijing Institute of Genomics, Chinese Academy of Science with project accession number (GSA: **CRA002536**) that are publicly accessible at http://bigd.big.ac.cn/gsa.

### SNV detection

The paired-end reads of WGS for each sample were aligned to the dog reference genome (CanFam3.1) using BWA-MEM (Version 0.7.17) with default parameters^45^. PCR (polymerase chain reaction) duplication reads were marked with MarkDuplicates in Picard tools (Version 2.15.0) and filtered with DuplicateReadFilter in GATK tools (Version 3.8-0). SNVs were called using MuTect2 for uniquely mapped reads with default parameters, where local *de-novo* assembly of the haplotypes was performed to increase the accuracy of mutation calling ^46,47^. The variant sites identified in normal dog populations that were considered as known SNPs (5,648,530 SNPs in total, ftp://ftp.ncbi.nih.gov/snp/organisms/dog_9615/) were filtered out. The candidate SNVs were further filtered using the following criteria: (1) the read depth of the SNV sites for both tumor and non-tumor sections were at least 10, (2) at least three reads supporting mutant alleles were required in the tumor sections, (3) only the SNVs supported by more than two reads in both forward and reverse strands were retained, (4) the mutant allele fraction of SNVs in tumor sample was greater than 0.05. The final SNVs were functionally annotated by both xenoRefGene using ANNOVA^48^ and the Variant Effect Predictor (VEP) (http://www.ensembl.org/vep).

### Calculation of Ka/Ks

For each gene or the genome-wide genes as a supergene, we have counted the number of synonymous (*s*) and non-synonymous substitutions (*n*) in coding regions, as well as the number of synonymous (*S*) and nonsynonymous sites (*N*) in total. Based on these, we calculated the ratio of non-synonymous and synonymous substitutions (*K*_*a*_/*K*_*s*_) using the following equation:

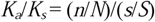

where, *s* for genes without synonymous substitutions was assumed to be 0.5 as suggested by Wang et al. (2011)^1,49,50^ to avoid calculation errors due to the few numbers of synonymous mutations detected in genes.

### Mutation spectrum and signature analysis

The MutationalPatterns package was utilized to extract and visualize mutational signatures in base substitution catalogues of tumor samples (https://github.com/UMCUGenetics/MutationalPatterns). Non-negative matrix factorization (NMF) was applied to extract *de novo* mutational signatures based on the mutation count matrix of 96 trinucleotide mutation classifications in canine mammary tumors ^51,52^. The number of signatures was determined according to the NMF quality measures of the cophenetic correlation coefficient and residual sum of squares (RSS) as suggested by Gaujoux and Seoighe (2010) ^53^. The optimal number should be chosen as the smallest number for which the cophenetic correlation coefficient starts decreasing and the number for which the plot of the RSS between the input matrix and its NMF estimate shows an inflection point. As shown in **Figure 3B**, the number of mutational signatures of 4 was selected.

The mutation catalogues (mutation types and counts) of the discovered signatures in canine mammary tumors were compared to those of the 30 standardized signatures in human cancer from the COSMIC (https://cancer.sanger.ac.uk/cosmic/signatures), and the cosine similarity between them was calculated with cos_sim_matrix^51^. Cosine similarity of 0.00 indicates completely different and similarity of 1.00 indicates a perfect match.

### Estimation of tumor purity and copy number alterations

We used the R package Sequenza to quantify cancer cellularity, ploidy, and chromosomal copy number alteration ^54^. The paired-end sequence reads of WGS for each sample were aligned to the dog reference genome (CanFam3.1) using BWA-MEM (Version 0.7.17) with default parameters^45^. The Pileup files of the tumor and normal specimens, as output by SAMtools, were processed by ‘sequenza-utils’ and binned by 1kb window size, calculating sequencing depth, homozygous and heterozygous positions in the normal sample, and variant alleles and allelic frequency in the tumor specimen. The *sequenza. the extract* was used to normalize the tumor versus normal depth ratios based on GC-content and perform allele-specific segmentation using the ‘copy number’ package. Based on the calculated B allele frequency and normalized depth ratio, the cellularity and ploidy were inferred from the point estimates with the maximum log posterior probability by fitting a Sequenza model using *sequenza*.*fit*. The allelic-specific copy numbers of chromosomal segments were further inferred according to the estimated cellularity and ploidy. Summary of the aberrant cell fraction and overall ploidy are provided in **Table S1**.

### Driver genes and regions analysis

We defined potential canine mammary cancer driver genes as previously described ^18^. The significance of broad and focal CNAs was assessed from the segmented data using the GISTIC 2.0 algorithm ^55^. We performed functional enrichment for these genes using DAVID ^56^.

### RNA Isolation and Total RNA Sequencing

Total RNA was extracted from 27 mammary gland tumors tissues using the RNeasy Mini Plus kit (Qiagen). Pulverization for sample homogenization was performed with liquid nitrogen before RNA isolation according to the manufacturer’s instructions. Cluster generation, followed by 2 × 100 cycle sequencing reads, separated by the paired-end turnaround, was performed on the instrument using HiSeq Rapid SBS Kit v2 (FC-402-4021) and HiSeq Rapid PE Cluster Kit v2 (PE-402-4002) (Illumina). Image analysis was performed using the HiSeq Control Software version 2.2.58. The raw data were processed, and base-calling was performed using the standard Illumina pipeline (CASAVA version 1.8.2 and RTA version 1.18.64). A summary of the statistics of the RNA sequencing data is listed in Table S8. RNA sequence data along genomic sequence data has been deposited in the genome sequence archive of Beijing Institute of Genomics, Chinese Academy of Science (http://gsa.big.ac.cn) with GSA accession number **CRA002536**.

### RNA-sequence analysis

For dog mammary gland tumors, we downloaded the RNA sequencing data of 4 pairs of normal and 9 tumor specimens from the NCBI SRA database with accession number **SRP024250** ^57^. Additionally, RNA sequencing is done for 27 canine mammary cancers are **Table S8**. The numbers of transcripts identified in the study were listed in Table S8. Paired-end reads of RNA-Seq were aligned to the dog reference genome (CanFam3.1) using the STAR (Version 2.6.0)^58^. Next, we calculated the TPM (transcripts per million) value for each gene using uniquely mapped reads with RSEM (RNA-Sequence by Expectation-Maximization, Version 1.3.0) software in a single-stranded mode^59^, which quantified the gene expression level accurately. The uniquely mapped paired (∼70-79%) was used to quantify a gene’s expression level by calculating its value using RSEM with the default parameters and the canine RefGene and Ensembl annotation downloaded from the NCBI and Ensembl genome site. Raw RSEM expected counts for all samples were normalized to the overall upper quartile. To identify the differentially expressed genes (DEGs) between different sample groups and calculate the significance level of expression differences, we performed the DESeq2 package (Version 1.14.0) with a negative binomial test based on the raw read depth ^60^.

Heatmaps were then generated by differentially expressed genes using their expected count via hierarchical clustering. Furthermore, gene functional annotation and enrichment analyses were achieved using the DAVID Functional Annotation tool (david.abcc.ncifcrf.gov)^56^.

### Construction of phylogenetic trees

We constructed a phylogenetic tree of the canine patients and tumors using the Maximum parsimony method in the PHYLIP package ^61^. After excluding SNVs located in LOH regions, the remaining SNVs were used to construct the phylogenetic tree. SNVs called from WGS was used for tree construction.

### Human cancer datasets

All VCF (Variant Call Format (Version 4.2)) files (called by Mutect2) for the human breast carcinoma (BRCA) were downloaded from the TCGA data coordinating centre (https://portal.gdc.cancer.gov/), each being reprocessed to eliminate known germline single nucleotide polymorphisms (SNP) present in the dbSNP database. Additionally, for a comprehensive understanding of somatic mutations in breast cancer across the genome, previously reported SNVs analyzed by using the WGS dataset, were downloaded from ftp://ftp.sanger.ac.uk/pub/cancer/Nik-ZainalEtAl-560BreastGenomes^20^. All variants coordinate mapped to the GRCh37, re-annotated using the ANNOVAR^48^, as it would have required a uniform variant calling and filtering workflow. RNA-seq dataset downloaded from publicly available cancer transcriptome data (TCGA and MET500 transcriptome sequencing), http://ualcan.path.uab.edu/index.html ^62^.

## Author contributions

SA, XL, and YH conceived and designed the experiments, SA and WD performed the DNA and RNA extraction, library construction, SA, LG, XB, and YH performed the data analysis, SA, YH and XL wrote the manuscript, D.L, G.B, Y.S, and YH provided and collected the samples.

## Acknowledgments

The authors delightedly acknowledge the Strategic Priority Research Program of the Chinese Academy of Sciences (XDB13040300), Youth Innovation Promotion Association, CAS-TWAS president’s fellowship of Ph.D., and Chinese Academy of Sciences for sponsoring this research.

## FIGURE LEGENDS

**Table 1. The 127 SMGs from 20 cellular processes in mammary cancer**

**Figure S1.**
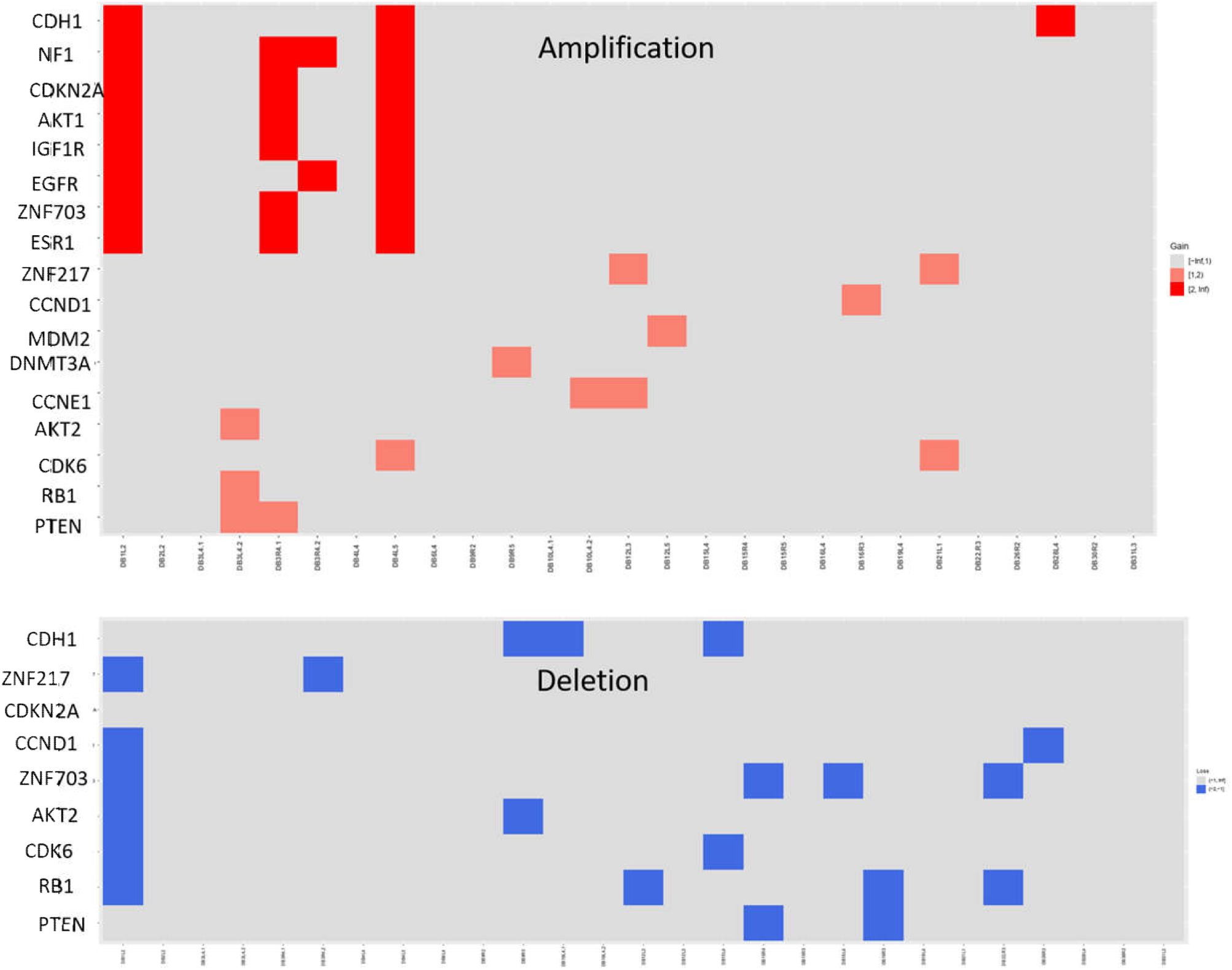
The heatmap showing the segmented copy-number profiles in Canine mammary cancer. The chromosomes are arranged vertically from top to bottom and samples are arranged from left to right. Red and blue represent gain and loss, respectively.

**Figure S2.**
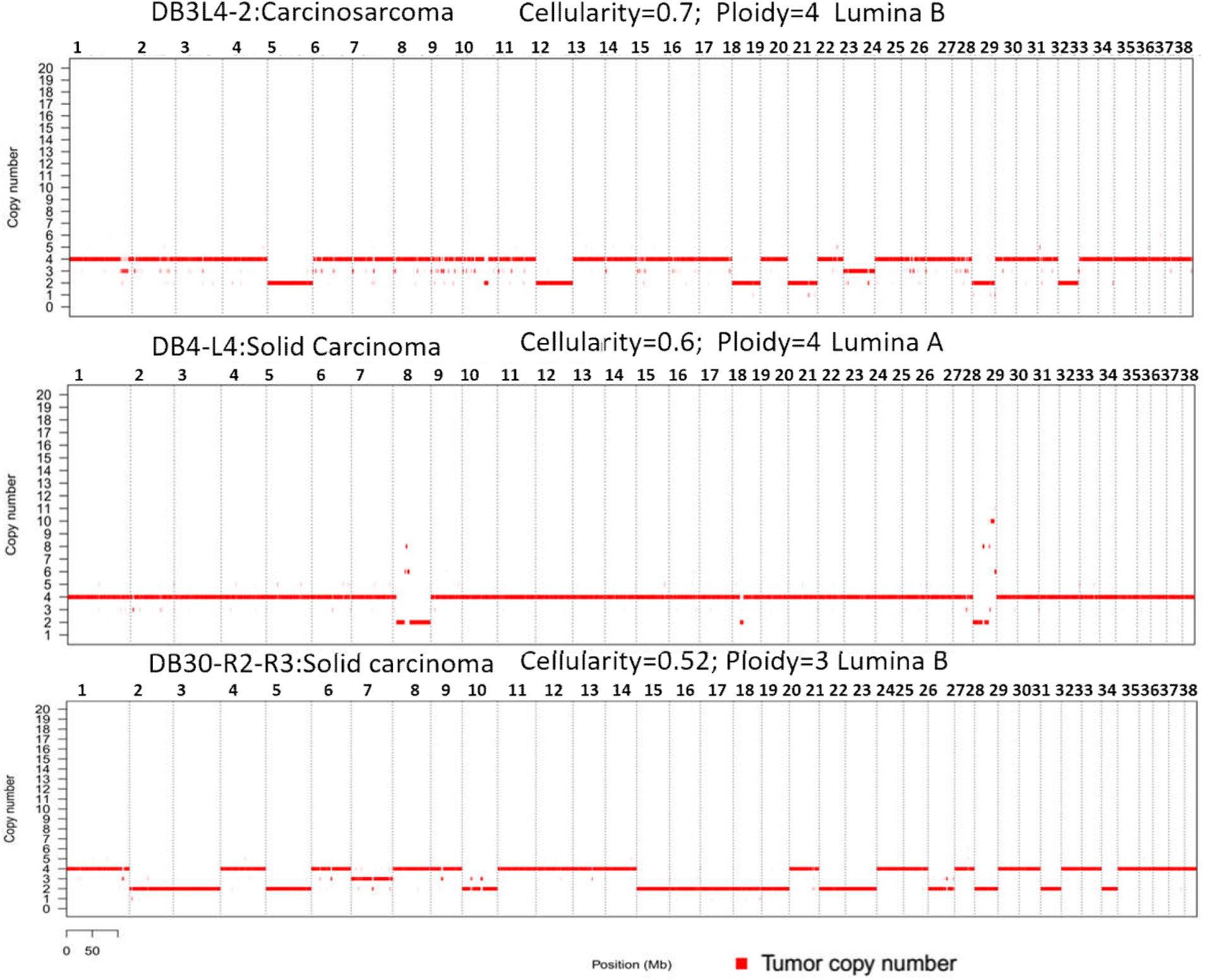
Tumor copy number profile of tetraploidy DB3L4-2, DB4-L4 and DB3-R2-R3. X-axis represents each chromosome, and Y-axis represents copy number in the tumor.

**Figure S3.**
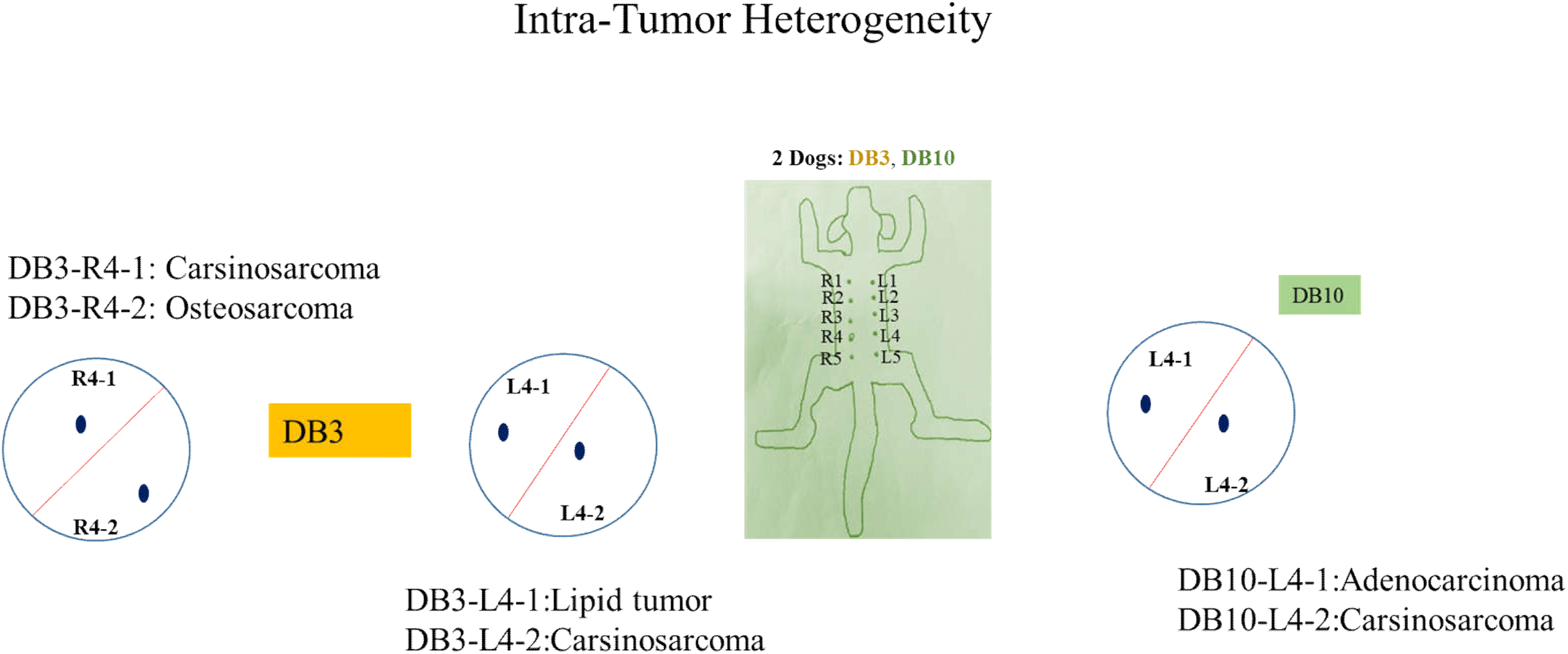
Sampling strategy of Dog 3 and Dog10 intratumor heterogeneity. The blue dots show one sample from each part within tumor. Each circle shows one tumor.

**Figure S4.**
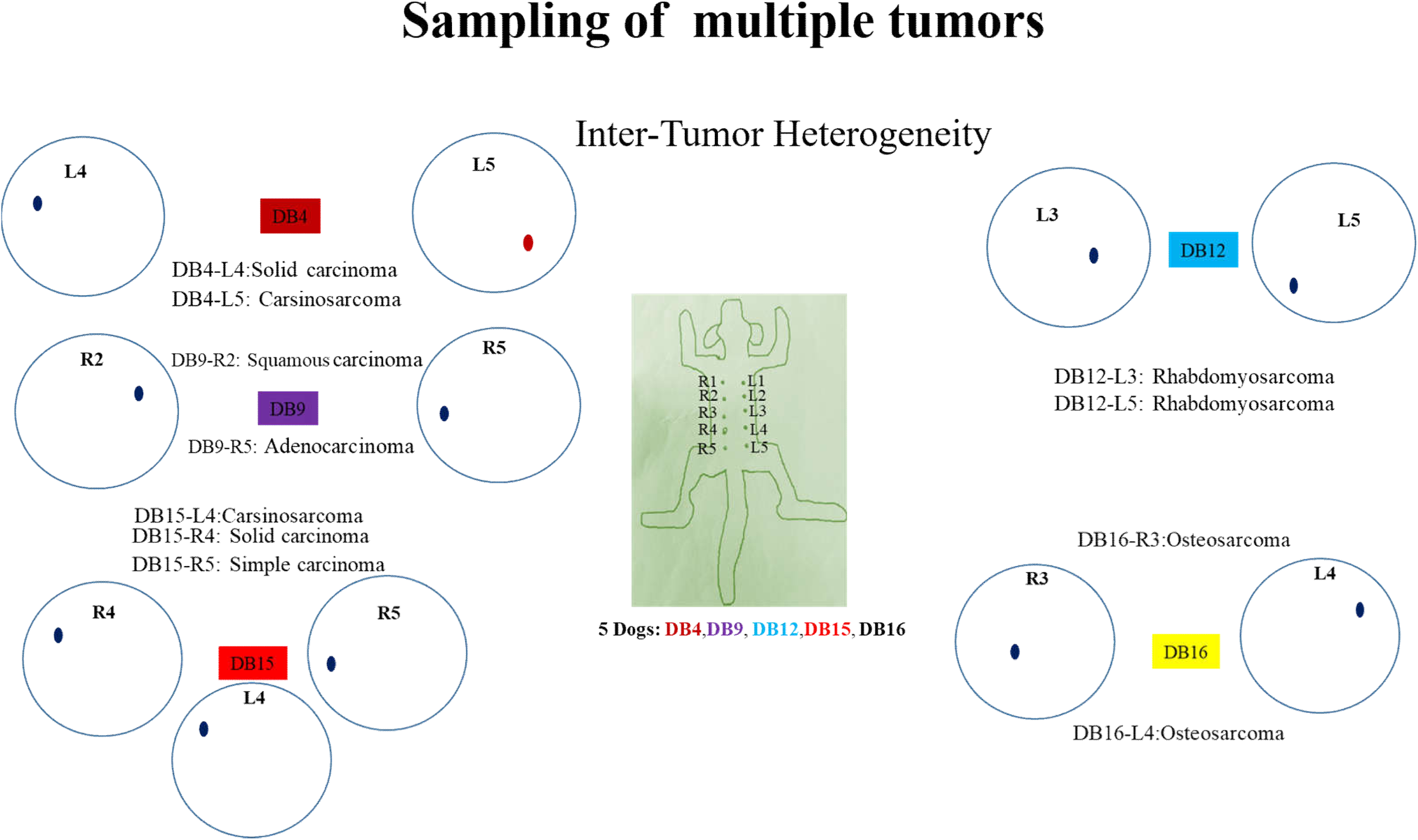
Sampling strategy of Dog 4, 9, 12, 15 and Dog16 inter-tumor heterogeneity. The blue dots show one sample from each tumor. Each circle shows one tumor.

**Figure S5.**
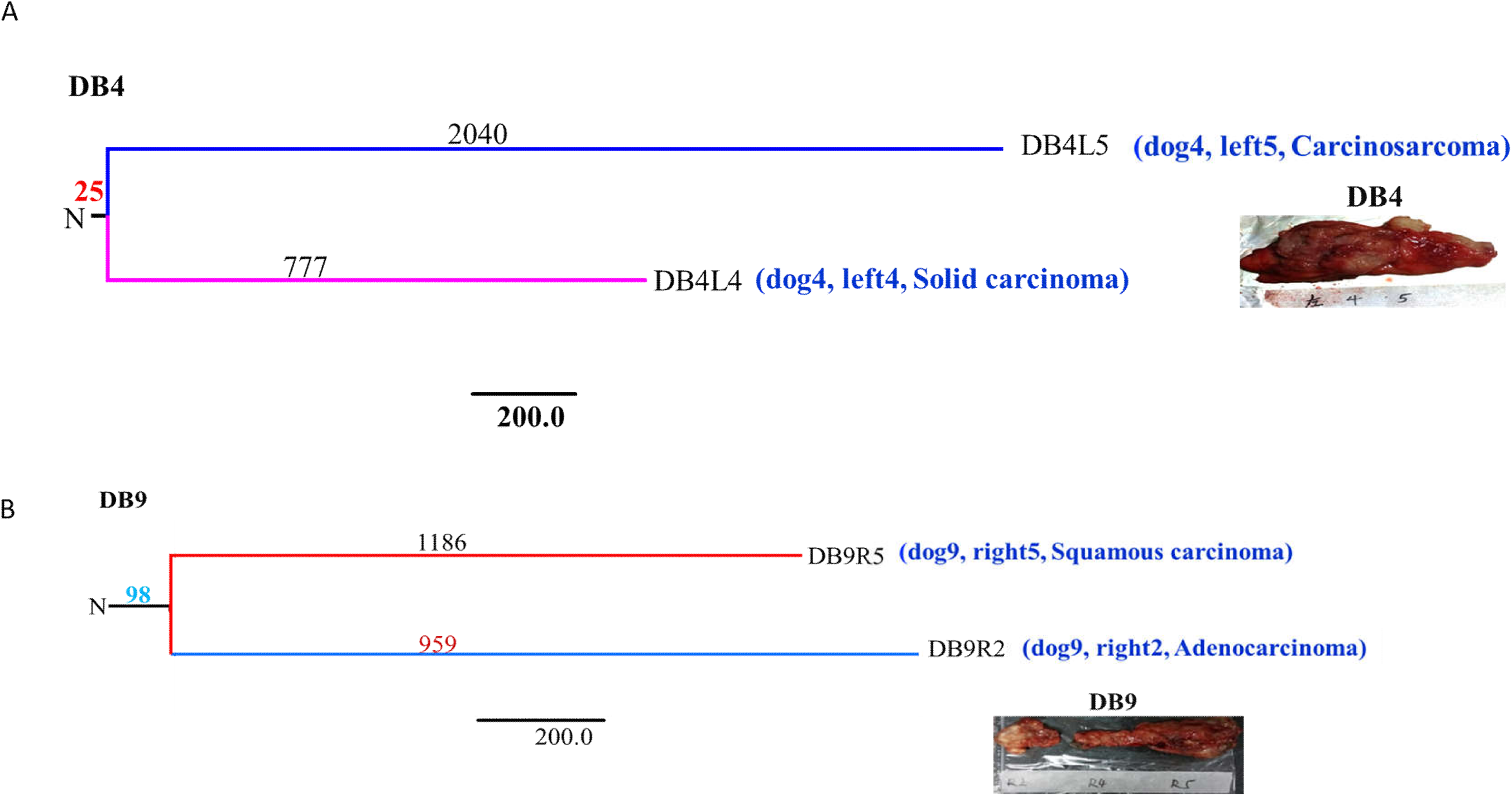
A. Phylogenetic relationship between the tumors in Dog 4 between four samples. The lengths of branches are proportional to mutation numbers. The tree was anchored using a DNA sequence from the normal tissue of a canine patient. **B. Phylogenetic relationship between two samples in Dog9**. The tree was anchored using a DNA sequence from the normal tissue of the canine patient. The lengths of branches are proportional to mutation numbers. The numbers correspond to the number of common mutations and specific mutations in the samples.

**Figure S6.**
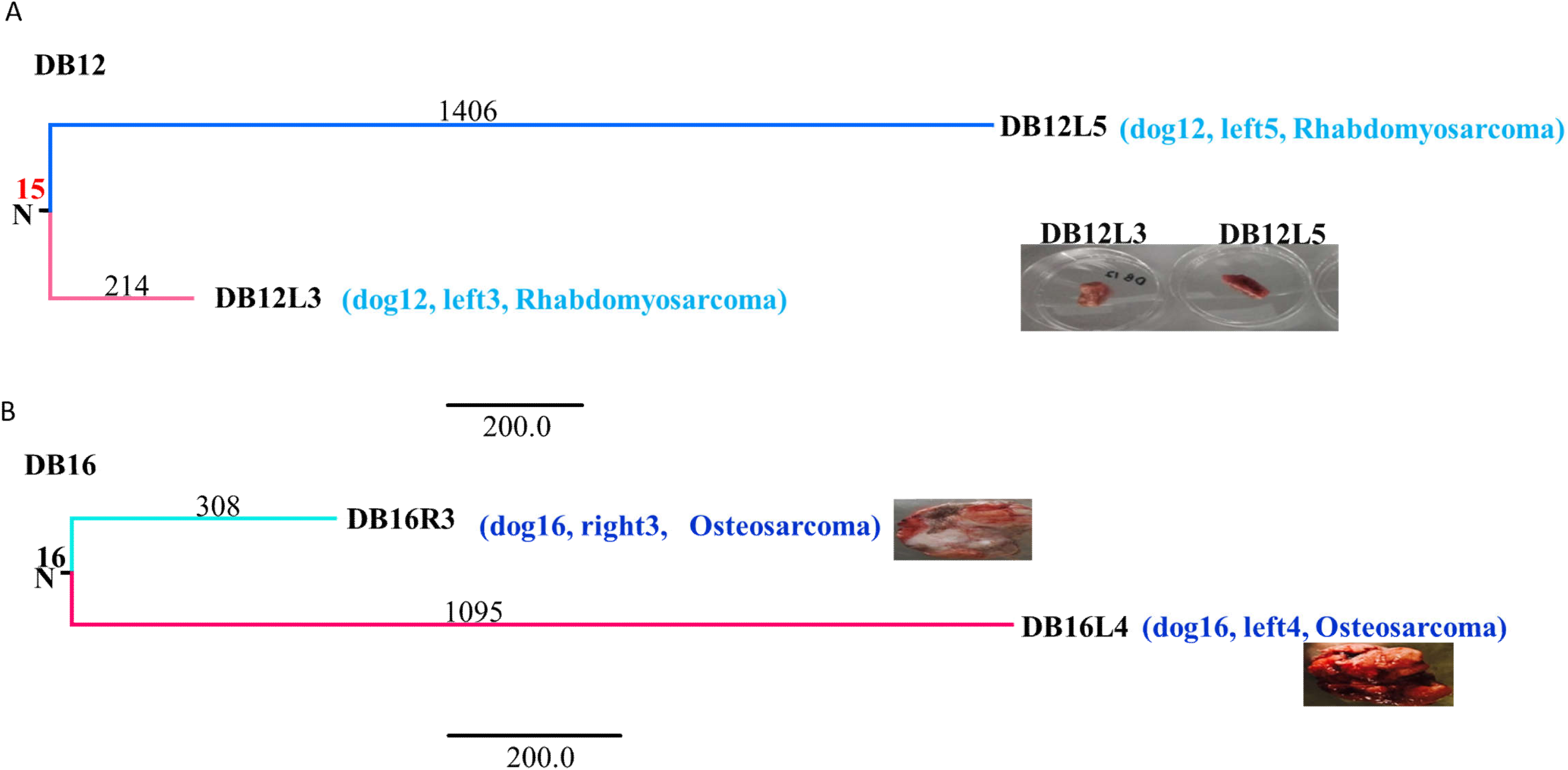
A. Phylogenetic relationship between two samples in Dog12. The lengths of branches are proportional to mutation numbers. The tree was anchored using a DNA sequence from the normal tissue of a canine patient. **B. Phylogenetic relationship between two samples in Dog16**. The tree was anchored using a DNA sequence from the normal tissue of the canine patient. The lengths of branches are proportional to mutation numbers. The numbers correspond to the number of common mutations and specific mutations in the samples.

## Tables Provided as Excel files

**Table S1**. Supplementary data for canine mammary case information, including histopathological description, ER, PR and HER status, age, breed, and sex. This table provides supplementary data for Figure 1.

**Table S2**. Supplementary data for the TNM staging system for canine mammary carcinomas.

**Table S3**. Supplementary data mutation discovery in canine mammary cancer and human breast cancer by WGS and WES, total SNVs listed across the genome.

**Table S4**. Supplementary data for mutated genes with indels, GO for indels. And SNVs analysis Fisher test.

**Table S5**. Supplementary data for the percentage of a mutated gene listed SNVs base substitution type occurrence in human and dog. The supporting data for Table 1.

**Table S6**. Supplementary data for somatic copy number variation from GISTIC 2.0 analysis.

**Table S7**. Supplementary data, GO for somatic copy number variation.

**Table S8**. Supplementary data for RNA-seq sequencing and mapping.

**Table S9**. Supplementary data for differentially expressed genes among canine carcinomas and Human breast cancer, including the gene list and p-value and log fold change among the genes.

**Table S10**. Supplementary data for GO of differentially expressed genes among canine carcinomas and Human breast cancer, including the gene list and enriched functions among the genes.

**Table S11**. Supplementary data for data for Ka/Ks for aging-related genes in mammary cancer in both species. The supporting data for Figure 7.

**Table S12**. Supplementary data for inter and intra tumor heterogeneity.

## Reference

1 Wu, C. I., Wang, H. Y., Ling, S. & Lu, X. The Ecology and Evolution of Cancer: The Ultra-Microevolutionary Process. Annu Rev Genet 50, 347–369, doi:10.1146/annurev-genet-112414-054842 (2016).

2 Ling, S. et al. Extremely high genetic diversity in a single tumor points to prevalence of non-Darwinian cell evolution. Proceedings of the National Academy of Sciences 112, E6496–6505, doi:10.1073/pnas.1519556112 (2015).

3 Sottoriva, A. et al. A Big Bang model of human colorectal tumor growth. Nature Genetics 47, 209–216, doi:10.1038/ng.3214 (2015).

4 Li, C. et al. A Direct Test of Selection in Cell Populations Using the Diversity in Gene Expression within Tumors. Mol Biol Evol 34, 1730–1742, doi:10.1093/molbev/msx115 (2017).

5 Wong, K. et al. Cross-species genomic landscape comparison of human mucosal melanoma with canine oral and equine melanoma. Nat Commun 10, 353, doi:10.1038/s41467-018-08081-1 (2019).

6 Wu, C.-I., Wang, H.-Y., Ling, S. & Lu, X. The Ecology and Evolution of Cancer: The Ultra-Microevolutionary Process. Annual Review of Genetics 50, 347–369, doi:10.1146/annurev-genet-112414-054842 (2016).

7 Ostrander, E. A., Dreger, D. L. & Evans, J. M. Canine Cancer Genomics: Lessons for Canine and Human Health. Annu Rev Anim Biosci 7, 449–472, doi:10.1146/annurev-animal-030117-014523 (2019).

8 Pinho, S. S., Carvalho, S., Cabral, J., Reis, C. A. & Gartner, F. Canine tumors: a spontaneous animal model of human carcinogenesis. Translational Research 159, 165–172, doi:10.1016/j.trsl.2011.11.005 (2012).

9 Liu, D. et al. Molecular homology and difference between spontaneous canine mammary cancer and human breast cancer. Cancer Res 74, 5045–5056, doi:10.1158/0008-5472.CAN-14-0392 (2014).

10 Dobson, J. M., Samuel, S., Milstein, H., Rogers, K. & Wood, J. L. Canine neoplasia in the UK: estimates of incidence rates from a population of insured dogs. Journal of Small Animal Practice 43, 240–246 (2002).

11 Medawar, P. B. An Unsolved Problem of Biology. (1952).

12 Williams, G. C. Pleiotropy, Natural Selection, and the Evolution of Senescence. Evolution Vol. 11, pp. 398–411 (1957).

13 Holliday, R. Aging is no longer an unsolved problem in biology. Annals of the New York Academy of Sciences 1067, 1–9, doi:10.1196/annals.1354.002 (2006).

14 Devarakonda, S., Morgensztern, D. & Govindan, R. Genomic alterations in lung adenocarcinoma. Lancet Oncol 16, e342–351, doi:10.1016/S1470-2045(15)00077-7 (2015).

15 Alexandrov, L. B. et al. Clock-like mutational processes in human somatic cells. Nature Genetics 47, 1402–1407, doi:10.1038/ng.3441 (2015).

16 Medawar, P. B. An unsolved problem of biology. 24 pp (Published for the College by H.K. Lewis, 1952).

17 Lee, K. H., Park, H. M., Son, K. H., Shin, T. J. & Cho, J. Y. Transcriptome Signatures of Canine Mammary Gland Tumors and Its Comparison to Human Breast Cancers. Cancers 10, doi:10.3390/cancers10090317 (2018).

18 Nik-Zainal, S. et al. Landscape of somatic mutations in 560 breast cancer whole-genome sequences. Nature 534, 47–54, doi:10.1038/nature17676 (2016).

19 Sakthikumar, S. et al. SETD2 is recurrently mutated in whole-exome sequenced canine osteosarcoma. Cancer Research, canres.3558.2017, doi:10.1158/0008-5472.can-17-3558 (2018).

20 Nik-Zainal, S. et al. Landscape of somatic mutations in 560 breast cancer whole-genome sequences. Nature 534, 47-+, doi:10.1038/nature17676 (2016).

21 Kandoth, C. et al. Mutational landscape and significance across 12 major cancer types. Nature 502, 333–339, doi:10.1038/nature12634 (2013).

22 Gkeka, P. et al. Investigating the structure and dynamics of the PIK3CA wild-type and H1047R oncogenic mutant. Plos Comput Biol 10, e1003895, doi:10.1371/journal.pcbi.1003895 (2014).

23 Martincorena, I. et al. Universal Patterns of Selection in Cancer and Somatic Tissues. Cell 173, 1823, doi:10.1016/j.cell.2018.06.001 (2018).

24 Van den Eynden, J., Basu, S. & Larsson, E. Somatic Mutation Patterns in Hemizygous Genomic Regions Unveil Purifying Selection during Tumor Evolution. PLoS Genet 12, e1006506, doi:10.1371/journal.pgen.1006506 (2016).

25 Li, Y. & de Magalhaes, J. P. Accelerated protein evolution analysis reveals genes and pathways associated with the evolution of mammalian longevity. Age (Dordr) 35, 301–314, doi:10.1007/s11357-011-9361-y (2013).

26 Wu, C. I., Wang, H. Y., Ling, S. & Lu, X. The Ecology and Evolution of Cancer: The Ultra-Microevolutionary Process. Annual Review of Genetics 50, 347–369, doi:10.1146/annurev-genet-112414-054842 (2016).

27 Carter, A. J. & Nguyen, A. Q. Antagonistic pleiotropy as a widespread mechanism for the maintenance of polymorphic disease alleles. BMC Medical Genetics 12, 160, doi:10.1186/1471-2350-12-160 (2011).

28 Liu, S. et al. Multi-omics Analysis of Primary Cell Culture Models Reveals Genetic and Epigenetic Basis of Intratumoral Phenotypic Diversity. Genomics Proteomics Bioinformatics 17, 576–589, doi:10.1016/j.gpb.2018.07.008 (2019).

29 Li, C. et al. A Direct Test of Selection in Cell Populations Using the Diversity in Gene Expression within Tumors. Molecular Biology and Evolution 34, 1730–1742, doi:10.1093/molbev/msx115 (2017).

30 Bedard, P. L., Hansen, A. R., Ratain, M. J. & Siu, L. L. Tumour heterogeneity in the clinic. Nature 501, 355–364, doi:10.1038/nature12627 (2013).

31 Tao, Y. et al. Further genetic diversification in multiple tumors and an evolutionary perspective on therapeutics. bioRxiv, 025429, doi:10.1101/025429 (2015).

32 Roche, J. The Epithelial-to-Mesenchymal Transition in Cancer. Cancers (Basel) 10, doi:10.3390/cancers10020052 (2018).

33 Sakthikumar, S. et al. <em>SETD2</em> Is Recurrently Mutated in Whole-Exome Sequenced Canine Osteosarcoma. Cancer Research 78, 3421–3431, doi:10.1158/0008-5472.can-17-3558 (2018).

34 Gavrilov, L. A. & Gavrilova, N. S. Evolutionary theories of aging and longevity. The Scientific World Journal 2, 339–356, doi:10.1100/tsw.2002.96 (2002).

35 Van den Eynden, J., Basu, S. & Larsson, E. Somatic Mutation Patterns in Hemizygous Genomic Regions Unveil Purifying Selection during Tumor Evolution. PLOS Genetics 12, e1006506, doi:10.1371/journal.pgen.1006506 (2016).

36 Crespi, B. J. & Summers, K. Positive selection in the evolution of cancer. Biol Rev 81, 407–424, doi:10.1017/S1464793106007056 (2006).

37 Flatt, T. & Partridge, L. Horizons in the evolution of aging. BMC Biology 16, 93, doi:10.1186/s12915-018-0562-z (2018).

38 Finch, C. E. Evolution in health and medicine Sackler colloquium: Evolution of the human lifespan and diseases of aging: roles of infection, inflammation, and nutrition. Proceedings of the National Academy of Sciences of the United States of America 107 Suppl 1, 1718–1724, doi:10.1073/pnas.0909606106 (2010).

39 Daniel Fabian, T. F. The Evolution of Aging. Nature Education Knowledge 3(10) (2011).

40 Wang, G. D. et al. The genomics of selection in dogs and the parallel evolution between dogs and humans. Nat Commun 4, 1860, doi:10.1038/ncomms2814 (2013).

41 Withrow, S. J., Vail, D. M. & Page, R. L. Withrow & MacEwen’s small animal clinical oncology. (2013).

42 Tavasoly, A., Golshahi, H., Rezaie, A. & Farhadi, M. Classification and grading of canine malignant mammary tumors. Vet Res Forum 4, 25–30 (2013).

43 Port Louis, L. R., Varshney, K. C. & Nair, M. G. An immunohistochemical study on the expression of sex steroid receptors in canine mammary tumors. ISRN Vet Sci 2012, 378607, doi:10.5402/2012/378607 (2012).

44 Cassali, G. et al. Consensus regarding the diagnosis, prognosis and treatment of canine mammary tumors: Benign mixed tumors, carcinomas in mixed tumors and carcinosarcomas. Vol. 10 (2017).

45 Li, H. & Durbin, R. Fast and accurate long-read alignment with Burrows-Wheeler transform. Bioinformatics 26, 589–595, doi:10.1093/bioinformatics/btp698 (2010).

46 Xu, C. A review of somatic single nucleotide variant calling algorithms for next-generation sequencing data. Comput Struct Biotechnol J 16, 15–24, doi:10.1016/j.csbj.2018.01.003 (2018).

47 Cibulskis, K. et al. Sensitive detection of somatic point mutations in impure and heterogeneous cancer samples. Nat Biotechnol 31, 213–219, doi:10.1038/nbt.2514 (2013).

48 Wang, K., Li, M. & Hakonarson, H. ANNOVAR: functional annotation of genetic variants from high-throughput sequencing data. Nucleic Acids Res 38, e164, doi:10.1093/nar/gkq603 (2010).

49 Ostrow, S. L., Barshir, R., DeGregori, J., Yeger-Lotem, E. & Hershberg, R. Cancer Evolution Is Associated with Pervasive Positive Selection on Globally Expressed Genes. Plos Genetics 10, doi:ARTN e1004239 10.1371/journal.pgen.1004239 (2014).

50 Zhou, Z. et al. Mutation-profile-based methods for understanding selection forces in cancer somatic mutations: a comparative analysis. Oncotarget 8, 58835–58846, doi:10.18632/oncotarget.19371 (2017).

51 Blokzijl, F., Janssen, R., van Boxtel, R. & Cuppen, E. MutationalPatterns: comprehensive genome-wide analysis of mutational processes. Genome Med 10, 33, doi:10.1186/s13073-018-0539-0 (2018).

52 Gaujoux, R. & Seoighe, C. A flexible R package for nonnegative matrix factorization. BMC Bioinformatics 11, 367, doi:10.1186/1471-2105-11-367 (2010).

53 Gaujoux, R. & Seoighe, C. A flexible R package for nonnegative matrix factorization. BMC Bioinformatics 11, 367, doi:10.1186/1471-2105-11-367 (2010).

54 Favero, F. et al. Sequenza: allele-specific copy number and mutation profiles from tumor sequencing data. Ann Oncol 26, 64–70, doi:10.1093/annonc/mdu479 (2015).

55 Mermel, C. H. et al. GISTIC2.0 facilitates sensitive and confident localization of the targets of focal somatic copy-number alteration in human cancers. Genome Biol 12, R41, doi:10.1186/gb-2011-12-4-r41 (2011).

56 Huang, D. W. et al. DAVID Bioinformatics Resources: expanded annotation database and novel algorithms to better extract biology from large gene lists. Nucleic Acids Research 35, W169–175, doi:10.1093/nar/gkm415 (2007).

57 Liu, D. et al. Molecular homology and difference between spontaneous canine mammary cancer and human breast cancer. Cancer Research 74, 5045–5056, doi:10.1158/0008-5472.CAN-14-0392 (2014).

58 Dobin, A. et al. STAR: ultrafast universal RNA-seq aligner. Bioinformatics 29, 15–21, doi:10.1093/bioinformatics/bts635 (2013).

59 Li, B. & Dewey, C. N. RSEM: accurate transcript quantification from RNA-Seq data with or without a reference genome. BMC Bioinformatics 12, 323, doi:10.1186/1471-2105-12-323 (2011).

60 Anders, S. & Huber, W. Differential expression analysis for sequence count data. Genome Biol 11, R106, doi:10.1186/gb-2010-11-10-r106 (2010).

61 Felsenstein, J. PHYLIP-Phylogeny Inference Package (Version 3.2). Cladistics 5, 164–166, doi:citeulike-article-id:2344765 (1989).

62 Chandrashekar, D. S. et al. UALCAN: A Portal for Facilitating Tumor Subgroup Gene Expression and Survival Analyses. Neoplasia 19, 649–658, doi:10.1016/j.neo.2017.05.002 (2017).

